# Cue-driven motor planning facilitates express visuomotor responses in human arm muscles

**DOI:** 10.1101/2022.04.02.486810

**Authors:** Samuele Contemori, Gerald E. Loeb, Brian D. Corneil, Guy Wallis, Timothy J. Carroll

**Affiliations:** Centre for Sensorimotor Performance, School of Human Movement and Nutrition Sciences, The University of Queensland, Brisbane, Australia; Department of Biomedical Engineering, University of Southern California, Los Angeles, California, USA; Department of Physiology and Pharmacology, Western University, London, Ontario, Canada; Department of Psychology, Western University, London, Ontario, Canada; Robarts Research Institute, London, Ontario, Canada

**Keywords:** motor preparation, stimulus-locked response, subcortical motor control, superior colliculus, reticular formation, stimulus predictability

## Abstract

Humans can produce “express” (∼100ms) arm muscle responses that are inflexibly locked in time and space to the presentation of a visual target, consistent with subcortical visuomotor transformations via the tecto-reticulo-spinal pathway. These express visuomotor responses are sensitive to explicit cue-driven expectations, but it is unclear at what stage of sensory-to-motor transformation such modulation occurs. Here, we recorded electromyographic activity from shoulder muscles as participants reached toward one of four virtual targets whose physical location was partially predictable from a symbolic cue. In an experiment in which targets could be veridically reached, express responses were inclusive of the biomechanical requirements for reaching the cued locations and not systematically modulated by cue validity. In a second experiment, movements were restricted to the horizontal plane so that the participants could perform only rightward or leftward reaches, irrespective of target position on the vertical axis. Express muscle responses were almost identical for targets that were validly cued in the horizontal direction, regardless of cue validity in the vertical dimension. Together, these findings suggest that the cue-induced enhancements of express responses are dominated by effects at the level of motor plans and not solely via facilitation of early visuospatial target processing. Notably, direct cortico-tectal and cortico-reticular projections exist that are well-placed to modulate pre-stimulus motor preparation state in subcortical circuits. Our results appear to reflect a neural mechanism by which contextually relevant motor plans can be stored within subcortical visuomotor nodes and rapidly released in response to compatible visual inputs.

**NEW & NOTEWORTHY:** Express arm muscle responses to suddenly appearing visual targets for reaching rapid have been attributed to the tecto-reticulo-spinal pathway in humans. We demonstrate that symbolic cues before target presentation can modulate such express arm muscle responses compatibly with the biomechanics of the cued reaching direction and the cue validity. This implies cortically mediated modulation of one or more sensorimotor transformation nodes of the subcortical express pathway.

## INTRODUCTION

Reaching for objects requires neural computations to evaluate the surrounding context and transform the sensory information into appropriate agonist\antagonist muscle responses to bring the hand to the target (1, 2, 3). In behavioural neuroscience, the time required by the brain to initiate a visually guided action is often inferred from the stimulus-to-movement delay (i.e. reaction time, RT; 4). The time that agonist and antagonist muscles take to produce a sufficient net internal joint torque to overcome the limb inertia and start the movement, however, is longer than the earliest stimulus-related electromyographic (EMG) muscle response. Therefore, earlier evidence of visuomotor processes at play during a target-directed reaching task can be provided by measuring EMG signals than RTs.

Interestingly, previous EMG work shown that humans can produce extremely fast stimulus-driven arm muscles responses that are inflexibly locked in space and time to visual stimuli (5-18), which were originally termed stimulus-locked responses (SLRs; 5). Specifically, the SLRs consistently encode the location of a visual stimulus within ∼100ms from its presentation, irrespective of the mechanical RT. The SLRs hence lack the typical trial-by-trial variability of volitional muscle response and ensuing RT that is due to multiple external and internal factors (e.g. number of choices, attention, motivation, expectation). This suggests that the visuomotor pathway for this specific class of short-latency stimulus-locked responses differs from longer-latency “movement-locked” muscle responses. Indeed, the neural pathway for SLRs has been hypothesized to parallel that of express saccades (19), which involves the superior colliculus and its downstream projection to the reticular formation (20-24). Consequently, we and others have referred to short-latency EMG responses observed within the established “SLR” time window as “express” arm muscles responses (16-18), and will use this terminology here.

Express arm muscle responses were recently shown to be modulated by temporal expectations about the stimulus (16), suggesting a top-down cortical modulation of the putative subcortical express pathway. In addition, Contemori et al. (17) showed that explicitly cueing the target location with a symbolic arrow-shaped cue promoted or impaired express muscle responses to cued or non-cued targets, respectively. This could reflect top-down influence on express muscle response via modulation of cue-driven *sensory* or *motor* processing. In the first case, the cue might direct attention to the expected target location in the visual field thus facilitating sensory-to-motor transformation of validly cued targets. In the second case, the cue might enhance pre-target motor preparation of the expected reach such that express muscle responses are facilitated when a compatible target appears. Here, we ran two experiments to dissociate these possible mechanisms underlying the cue-induced modulations of express arm muscle responses in humans. To this aim, we implemented and validated a novel computational method to estimate onset times of express muscle responses in individual trials.

In the first experiment, participants reached toward one of four potential targets that were projected virtually in the horizontal plane, thus requiring different mechanical contributions from the muscles we tested to move the arm toward the four target locations. Specifically, the target could appear above or below a fixation spot either to the left or right of fixation. Trial-by-trial, the target location was partially predictable from a symbolic arrow-shaped cue oriented toward one of the four possible target locations such that its validity could be modulated on both right\left and top\bottom axes. If the cue modulates early visuospatial processing, then express muscle responses should be facilitated and impaired by valid and invalid cues, respectively. Although express muscle responses were influenced by the right\left cue validity, the validity effects of the top\bottom component of the cue were mixed. Indeed, express muscle responses were modulated consistently with the mechanical output required from the recorded muscle to reach toward the cued direction, irrespective of the actual top\bottom target location. This suggests that the cue-induced modulations of express responses act, at least partly, at the level of motor plans along the putative subcortical express pathway, rather than solely via facilitation of early sensorimotor transformations at cued spatial location.

We next tested whether cueing the likely target location influences express visuomotor behaviour via purely visuospatial mechanisms, in addition to any effect on motor preparation. We used a target paradigm similar to that of the first experiment, but here the targets were projected virtually in the vertical plane while reaching movements were restricted to the horizontal plane. This allowed the participants to use the right\left cue orientation to prepare a reach that would be appropriate for either the top or the bottom target. Thus the top\bottom cue orientation was relevant only to identify the location in the visual field at which the visual stimulus was expected to appear. We reasoned that if facilitation of visuospatial target processing contributes strongly to cue-enhanced express behaviour, then validly cueing the vertical location of the target should facilitate express responses. In contrast, we observed almost identical express arm muscle responses for targets that were compatible with the cued reaching direction, regardless of cue validity in the vertical dimension.

Overall, our results appear to reflect a contribution of top-down motor preparation to express arm muscle responses, such that there is rapid integration of motor plans for anticipated movements with emerging visual inputs along the putative subcortical express pathway. Such an arrangement could facilitate rapid release of prepared motor actions in response to compatible visual stimuli.

## MATERIALS AND METHODS

### Participants

Fifteen adults participated in the first experiment (3 females; mean age: 29.3±7.3 years), and eleven of them also participated in the second experiment that had a total sample of sixteen adults (2 females; mean age: 28.9±7.7 years). All participants were right-handed, had normal or corrected-to-normal vision, and reported no current neurological or musculoskeletal disorders. They provided informed consent and were free to withdraw from the experiment at any time. All procedures were approved by the University of Queensland Medical Research Ethics Committee (Brisbane, Australia) and conformed to the Declaration of Helsinki.

### Task design and experimental set-up

#### Task design

We used an *emerging moving* target paradigm (Figure 1A) that has proven effective to facilitate the express visuomotor arm muscle responses (14-18). In both experiments, the target was a filled black (∼0.3 cd/m^2^) circle of ∼2dva in diameter presented against a light grey background (∼140 cd/m^2^). This created a high target-to-background contrast necessary to facilitate the generation of express arm muscle responses (6, 15). The luminance was measured with a colorimeter (Cambridge Research System ColorCAL MKII). A photodiode was attached to the left bottom corner of the monitor to detect a secondary light that was presented coincidentally with the time of appearance of the real target. This allowed us to index the time point at which the stimulus was physically detectable.

**Figure 1:**
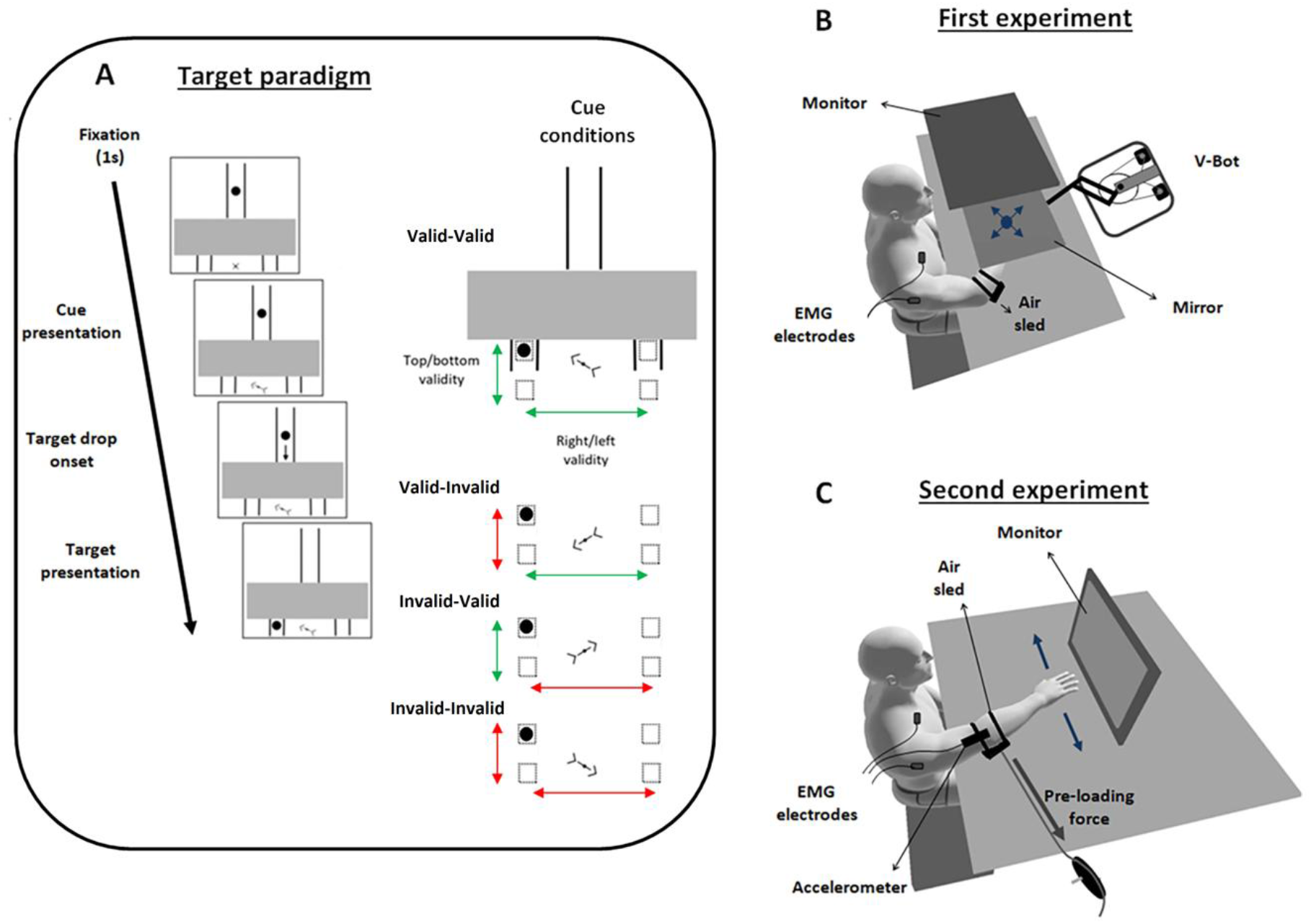
A, schematic diagram of the timeline of events in the target paradigm; a zoomed view of the paradigm and cue attributes are shown in the top left corner (the dashed boxes indicate the four possible locations of the target). After one second of fixation, the central “X” sign for fixation was substituted by an arrow cue pointing validly or invalidly toward the target location. After ∼700ms from cue presentation, the target started dropping from the stem of the track at constant velocity of ∼35dva/s until it passed behind the barrier (i.e. occlusion epoch) for ∼480ms, and re-appeared underneath it at ∼640ms from the its movement onset time (17). The target appeared transiently by making one single flash of ∼8ms of duration. The cue had two levels of validity: right/left and top/bottom (the cue validity and invalidity are shown with green and red arrows, respectively). The right column of panels shows the four cue-orientation variations relative to the target that appeared to the left and just beneath the barrier (i.e. top-left target). B, first experiment **e**xperimental setup. Participants’ hand positions were virtually represented via a cursor (blue dot) displayed on the monitor and projected into the (horizontal) plane of hand motion via a mirror. The head position was stabilized by a forehead rest (not shown here). The blue arrows represent the four reaching directions to address each of the four possible targets. C, second experiment experimental setup. In both experiments, participants were seated and began with their dominant (right) hand aligned with the fixation spot (“X” sign beneath the barrier, see panel A) and moved it toward a target that appeared beneath the barrier. Head position was stabilized by chin and forehead rests (not shown here). The blue arrows represent the rightward and leftward horizontal upper limb movements that participants could execute to address the location of one of the four possible targets (i.e. two possible movements for four possible targets).

The participants performed visually guided reaches toward the targets whose location was partially predictable from the orientation of a symbolic arrow-shaped cue (Figure 1A). The cue-target onset asynchrony (CTOA) was >1 second to ensure unambiguous interpretation of the arrow orientation. For both experiments, the target was constrained to fall within a track, which was shaped as an inverted diapason, until it passed behind a visual barrier that occluded the junction point at which the target randomly deviated left or right (16, 17). The target reappeared transiently (one single flash of ∼8ms of duration) at one of four different locations underneath the barrier: (i) to the right, just beneath the barrier (i.e. *top-right* location); (ii) to the right, closer to the bottom of the monitor (*bottom-right* location); (iii) to the left, just beneath the barrier (i.e. *top-left* location); (iv) to the left, closer to the bottom of the monitor (*bottom-left* location).

To start the trial, the participants were instructed to align their right hand (or the cursor in the first experiment; Figure 1B) and the gaze at a central “X” sign underneath the barrier (Figure 1A), and to stare at it for 1second. After the fixation period, the central fixation spot was changed to a symbolic inclined arrow oriented toward one of the four possible target locations. Note that symbolic meaning was derived from the orientation and shape of the cue, whereas its physical position was uninformative for the future target location. The cue informed the subjects about the likely right/left and top/bottom locations of the target, thus providing two levels of cue validity. There were four different cue conditions (right column of panels in Figure 1A): (I) ‘Valid-Valid’, when both the right/left and top/bottom target locations were validly cued; (II) ‘Valid-Invalid’, when only the right/left target location was validly cued; (III) ‘Invalid-Valid’, when only the top/bottom target location was validly cued; (IV) ‘Invalid-Invalid’, when neither the right/left nor the top/bottom target locations were validly cued.

#### Experiment 1: experimental set-up

Here we tested cue-induced modulations of express visuomotor responses when the four target locations could be reached via distinct and veridical movements. To this aim, we used a two-dimensional planar robotic manipulandum (the vBOT; 25). The target was displayed on a LCD computer monitor (120Hz refresh rate; 8.33ms/refresh cycle) mounted above the vBOT handle and projected to the participant via a mirror (Figure 1B). The stimuli were created in Microsoft Visual C++ (Version 14.0, Microsoft Visual Studio 2005) using the Graphic toolbox. The handle position was virtually represented by a blue cursor (∼1.2dva) whose apparent position coincided with actual hand position in the plane of the limb (i.e. top and bottom targets were physically distinguished by their depth relative to the body). The targets were located at ∼60°, ∼120°, ∼240° and ∼300° around the fixation spot (distance between top and bottom targets: ∼5.5cm; distance between right and left targets at the same depth: ∼10cm) and had equal eccentricity of ∼10dva from the fixation spot. A constant rightward load of ∼5N was applied to preload the shoulder transverse flexor muscles, including the clavicular head of pectoralis major muscle, and a custom-built air sled was positioned under the right elbow to minimize sliding friction (16, 17). The participants had to gaze at the fixation spot until the target reappeared from behind the barrier (a “fixation” error was shown if this condition was not met and the trial was reset), and then to start moving as rapidly as possible toward the target. Horizontal gaze-on-fixation was checked on-line with bitemporal, direct current electrooculography (EOG) sampled at 1 kHz. Each participant completed 15 blocks of 80 reaches/block (20 for each of the 4 target locations), with each block consisting of 56 Valid-Valid, 8 Valid-Invalid, 8 Invalid-Valid and 8 Invalid-Invalid cue trials, randomly intermingled (see the ‘Task design’ section and Figure 1A for further details). Therefore, the arrow validly cued the target location on 70% of the trials.

#### Experiment 2: experimental set-up

Here we tested the influence of prior information on express visuomotor responses toward the same four targets (Figure 1A) but the arm could move only horizontally and was unable to match their vertical locations. We used an experimental set-up previously described by Contemori et al. (16, 17) and illustrated in figure 1C. The participant executed right (extensor-ward) or left (flexor-ward) horizontal upper limb movements in response to targets presented at the four possible locations that were displayed on a LCD monitor (120Hz refresh rate; 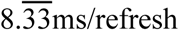 cycle) positioned vertically ∼57cm in front of the participants (distance between top and bottom targets: ∼8cm; distance between right and left targets: ∼15cm). Again, we preloaded (∼5N) the shoulder transverse flexor muscles and positioned a custom-built air sled under the right elbow to reduce movement friction. The stimuli were created in Matlab (version R2014b, TheMathWorks, Inc., Natick, Massachusetts, United States) using the Psychophysics toolbox (26, 27). The participants were instructed not to move their eyes from the fixation spot until the target reappeared from behind the barrier, and to reach as fast as possible toward the target. If the fixation condition was not met, the participants received an error message and the trial was reset. Gaze-on-fixation was checked on-line with an EyeLink 1000 plus tower-mounted eye tracker device (SR Research Ltd., Ontario, Canada), at a sampling rate of 1 kHz. Each participant completed 10 blocks of 68 reaches/block (17 for each of the 4 target locations) comprising: 40 Valid-Valid cues; 12 Valid-Invalid cues; 8 Invalid-Valid cues; 8 Invalid-Invalid cues. Therefore, the arrow cued the right\left target location with ∼75% validity, whereas the top\bottom location was cued with ∼75% validity for valid right\left cue conditions and 50% validity for invalid right\left cue conditions. The four cue conditions were randomly intermingled within each block.

### Data recording and analysis

#### Kinematic data

For the first experiment, we reconstructed the hand movements (Figure 2A) by using the vBOT handle kinematic data sampled at 1kHz. An error message was shown online if the cursor left the starting position within 130ms from the stimulus presentation. This RT cut-off was adopted because 130ms was recently shown to be the critical time to prepare a target-directed response (28). The RT for analysis was computed offline by identifying the first time point at which the radial (rho) hand velocity exceeded the baseline mean velocity (i.e. average velocity recorded from 100ms before the target presentation to the stimulus onset time) by more than five standard deviations (Figure 2B). Trials with RT<140ms (∼9%) and >500ms (<1%) were excluded from offline analysis. To determine the correct (i.e. target-directed) reaches, we recorded the *X* and *Y* hand position at the RT and when the hand reached 75% of the peak velocity (Figure 2C). We then computed the angle of a line passing between these hand positions to define the initial movement direction (Figure 2D). If the initial movement was directed within the quadrant of the visual field containing the target, we concluded that the movement was correct, otherwise the movement was classified as incorrect and not further analysed.

**Figure 2:**
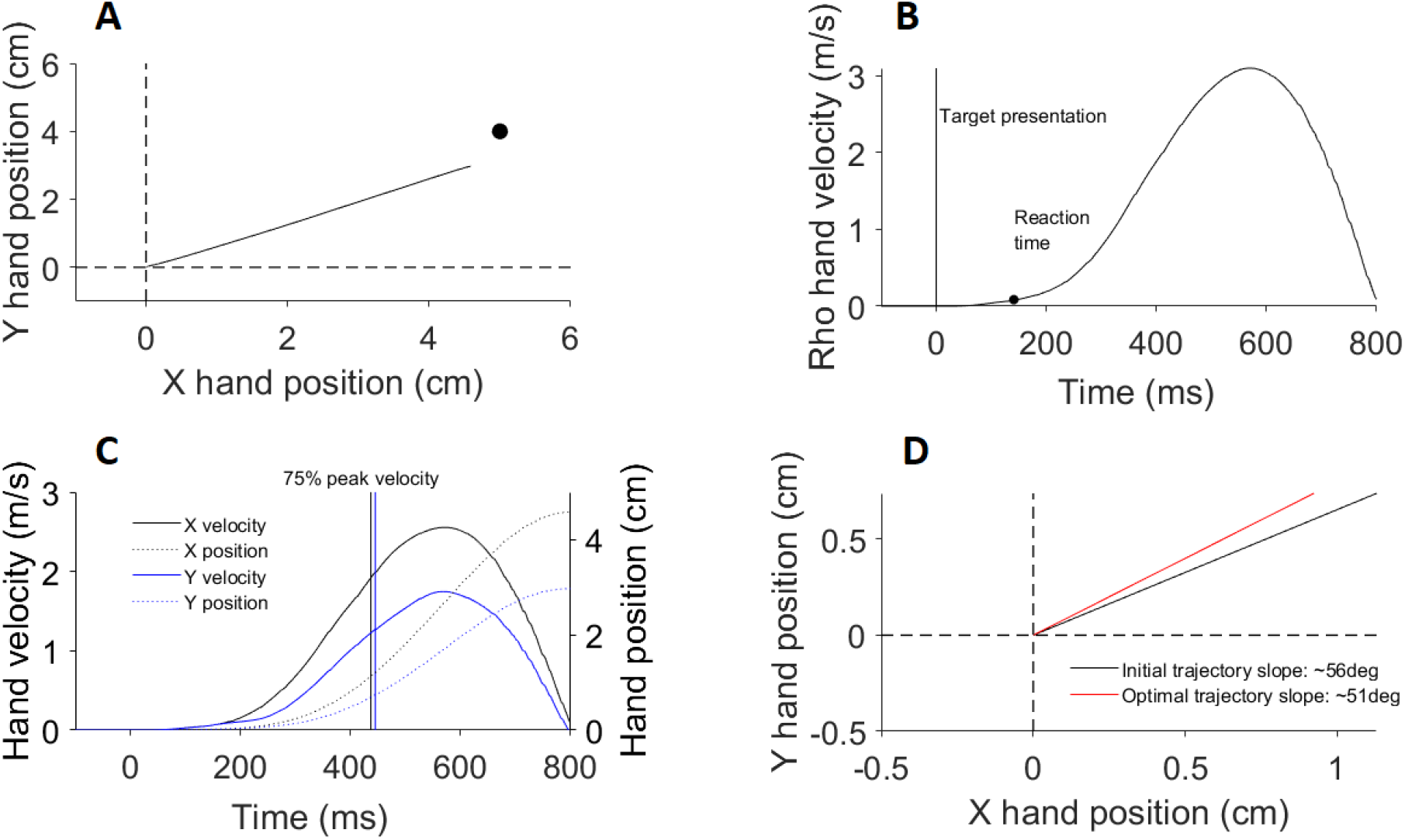
Kinematic analysis procedures for an exemplar valid cue trial. The movement was directed toward the target that appeared at the right of the fixation spot and just beneath the barrier (i.e. top-right target; see Figure 1A). A, hand reaching trajectory in the medio-lateral (X) and backward-forward (Y) axes from the [0,0] “home” position toward the target location (black dot). B, profile of the radial (rho) hand velocity used to index the reaction time of the movement after the target presentation. C, profiles of the hand velocity (solid lines) and trajectory (dotted lines) in the medio-lateral (X; black lines) and backward-forward (Y; blue lines) directions. The hand position coordinates at the reaction time and at 75% of the peak velocity in the X and Y axes (vertical lines) were used to determine the initial trajectory direction and the trial correctness. This trial was correct because the initial reach was directed within the quadrant of the visual field containing the target (see panel A). D, comparison between the optimal (∼56deg; red line) and actual (∼51deg; black line) initial hand trajectories. In this example, the initial trajectory error was +5deg indicating that the initial movement was directed slightly more horizontally than the optimal target trajectory.

To test whether the target-directed reach was modulated by the cue, we computed the angle error between the actual initial hand-to-target trajectory (see above) and the “optimal” hand-to-target trajectory. The optimal trajectory was defined by computing the angle of the line passing from the hand position at the RT and the target position. Note that the target position was corrected relative to the hand position at the RT, which on average was ∼0.5cm away from the “home” position. For each of the four target locations, negative angle errors indicate that the initial movement was further away from the horizontal midline between targets than the optimal target trajectory. By contrast, positive angle errors indicate that the initial reaching was closer to the horizontal midline between targets than the optimal target trajectory.

For the second experiment, the arm motion was monitored by a three-axis accelerometer (Dytran Instruments, Chatsworth, CA) sampled at 2kHz with a 16-bit analog-digital converter (USB-6343-BNC DAQ, National Instruments, Austin, TX, USA). Data synchronization was guaranteed on each trial by starting the recording at the frame at which the target started moving toward the barrier. We monitored the RT online (see 17 for details) and sent an error message if the participants moved before the target onset time or responded in <130ms from target presentation. The accelerometer signal was also used for offline RT computation and for the identification of correct (i.e. target-directed) responses (see 16 and 17 for details). Consistent with the first experiment, trials with RT<140ms (∼10%) and >500ms (<1%) were once again excluded during offline data analysis.

For both experiments, the kinematic data were averaged across the left and right directions to limit biases related to the preload. This was done separately for the top and bottom targets (i.e. kinematic data were pooled for right-up and left-up targets, and for right-down and left-down targets).

#### EMG data

Surface EMG activity was recorded from the clavicular head of the right pectoralis muscle (PMch) and the posterior head of the right deltoid muscle (PD) with double-differential surface electrodes (Delsys Inc. Bagnoli-8 system, Boston, MA, USA). Before the start of recording, we checked the quality of the EMG signal with an oscilloscope by asking the participants to flex (PMch activation-PD inhibition) and extend (PMch inhibition-PD activation) the shoulder in the transverse plane. The sEMG signals were amplified by 1,000, filtered with a 20–450 Hz bandwidth filter by the “Delsys Bagnoli-8 Main Amplifier Unit” and sampled at 2 kHz using a 16-bit analog-digital converter (USB-6343-BNC DAQ device, National Instruments, Austin, TX). The sEMG data were then down-sampled to 1 kHz and full-wave rectified offline without further filtering.

We and others have previously adopted a time-series *receiver operator characteristic* (ROC) analysis to determine the earliest stimulus-related muscle response (5, 16, 17). This analysis, however, is sensitive to the muscle response amplitude relative to the background signal-to-noise ratio, which is influenced by the sample size. Critically, Cross et al. (29) reported non-physiological short-latency muscle responses (<60ms) to visual perturbation of the cursor position when the ROC analysis was run on ∼10 trials. This calls into question the reliability of the ROC analysis for small data samples. In our experiments, relatively few trials (∼25) were run in the invalid cue conditions and trials with reaction times <140ms or >500ms and incorrect movements were excluded (see previous sections). Therefore, we developed a single-trial analysis method to extract the muscle response onset time from each correct trial. This analysis is described below and illustrated in figure 3. The ROC analysis was run, however, on the cue conditions with a large number of trials (i.e. Valid-Valid cue conditions; see previous section) to test the statistical contrasts between the two methods (Supplementary Materials and Methods; all Supplementary Material is available at https://osf.io/75jt2/?view_only=8cc72393af3642ac8628528325ccdf74).

**Figure 3:**
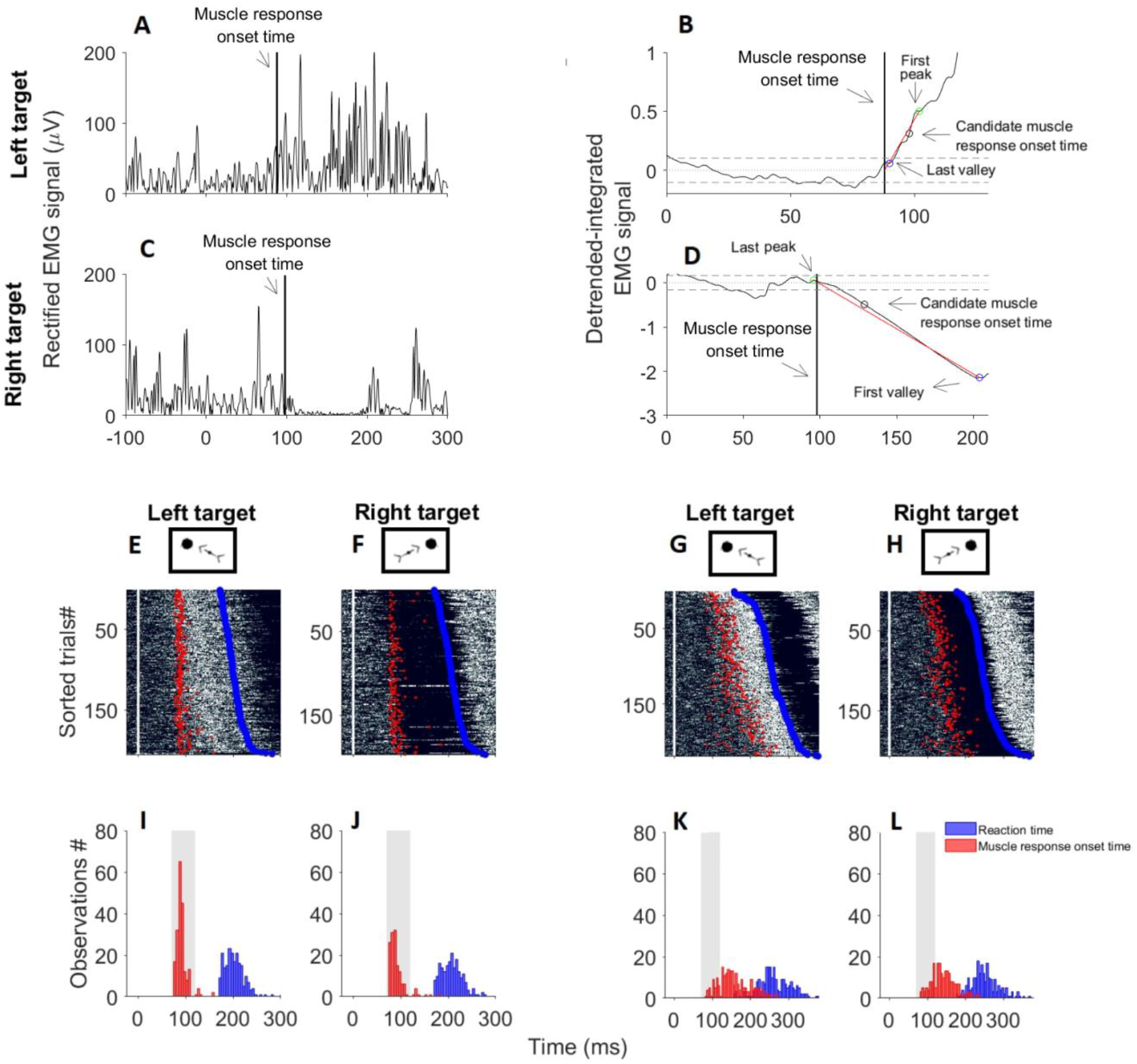
*Detrended-integrated* signal method to extrapolate the earliest stimulus-driven EMG response from each trial. The first two lines of panels show the procedures for an exemplar left target trial and an exemplar right target trial that were executed toward validly cued top targets. For both target locations, the rectified EMG activity is shown (panels A and C) as is a zoomed view of the detrended-integrated EMG signals (panels B and D). In these panels, the black unfilled scatter indicates the time at which the detrended-integrated signal diverges from background (dotted grey line) by more (B) or less (D) than five standard deviations (dashed grey lines). This point in time represents the candidate onset time of the express visuomotor muscle response. The green and blue unfilled scatters represent the boundaries of the time-window of data from which we computed the linear trendline (red line) that was used to index the point in time the linear trendline intercepted the background value of the detrended-integrated signal (solid vertical line in panels A-D), which represents the earliest muscle response initiation time. In these examples, the linear trendlines intercepted the background value of the detrended-integrated signal within 70-120ms (88ms for the left target; 98ms for the right target) after the target presentation and, thereby these trials were classified as express visuomotor muscle responses. Rasters of rectified surface EMG activity from individual trials are shown in panels E-H (brighter white colours indicate greater EMG activity). The white vertical line at 0ms indicates the target presentation time, the muscle response initiation time is represented with a red scatter and the blue scatters indicate the reaction time. Panels I-L show the distribution of muscle response onset time (red histograms) and reaction time (blue histograms), and the grey path indicates the time window in which an express muscle response is expected (70-120ms from target onset). The exemplar data showed in panels E and F are representative of an express muscle response producer because the earliest muscle responses appear as a vertical band of either muscle activations (E) or inhibitions (F) that is time locked ∼100 ms to the stimulus onset time, regardless of the movement onset time. Consistently, the distribution of the muscle response onset time is mostly enclosed within the 50ms express response time window (grey path in panels I and J) and does not match the spread of volitional movement onset times. By contrast, panels G and H represent a non-express muscle response because the muscle response onset time and reaction time are similarly distributed (panels K and L).

To index the earliest stimulus-related muscle response, we first subtracted the background activity value (i.e. average rectified EMG signal recorded from 100ms before to 70ms after the stimulus presentation) from the entire EMG signal. We then computed the integral of the EMG trace for each millisecond recorded between 100ms before and 300ms after the target onset time (Figure 3A-D). To remove the rising trend of the integrated signal, we *detrended* the signal by subtracting the linear regression function of the background period from the entire 400ms analysis window. We then computed the average and standard deviation values of the detrended-integrated signal in the background period, and indexed the candidate muscle response onset time as the first time the detrended-integrated signal exceeded the background value by more (i.e. earliest muscle activation), or less (i.e. earliest muscle inhibition), than five standard deviations (Figure 3B and D). Importantly, the occurrence of false-positive express muscle responses (i.e. candidate onset times earlier than 70ms after the target presentation) was lower than 5% with this threshold. Critically, the candidate response onset time is sensitive to the amplitude of the stimulus-driven response relative to the background activity and does not exactly correspond to the actual initial deviation of the EMG signal from background. To find this point in time, we ran a linear regression analysis around the candidate onset time of the stimulus-driven muscle response. Specifically, if the signal at the candidate onset time was higher than background, we extrapolated the linear trendline from the values enclosed between the last valley before and the first peak after the candidate onset time (Figure 3B). By contrast, if the signal at the candidate onset time was lower than background, the linear trendline was computed from the values enclosed between the last peak before and the first valley after the candidate onset time (Figure 3D). Finally, we defined the muscle response onset time as the point in time the linear trendline intercepted the background value of the detrended-integrated signal (vertical line in Figures 3B and D). Consistent with previous work (7, 16, 17), we defined the muscle response as “express” if it was initiated within 70-120ms after the target presentation. This method will be subsequently referred to as the *detrended-integrated* signal method.

Express muscle responses appear as a column of muscle activation (Figure 3E) or inhibition (Figure 3F) at ∼100ms after the stimulus onset time that does not co-vary with the voluntary movement initiation. In the absence of express responses, the earliest EMG responses (activation or inhibition) occur at times that co-vary with the RT (Figures 3G-H and K-L). To produce an objective measure for this distinction, we binned “fast” and “slow” trials according to the median RT value. We then associated the average response initiation time of the fast and slow trials with the average RT of the corresponding trials and we fitted a line to the data to test if the muscle response onset time did not co-vary with the RT. Specifically, we classified a data set as positive for an express muscle response if the slope of the line was >67.5 (5, 6, 16, 17). If express responses were positively detected for a data set, we computed the average muscle response initiation time and express response prevalence (%) across the individual trials within the data set with EMG onset times within the express muscle response window. We also quantified the express response magnitude by computing the average EMG activity recorded in the 10ms subsequent to the response initiation time for each rightward and leftward trial exhibiting an express muscle response. We then averaged this metric across the express response trials and computed the difference between the left and right targets. Note that these procedures were run separately for the top targets and the bottom targets to test the contrast between targets at comparable vertical locations.

#### Statistical analysis

Repeated measures ANOVA analyses with Bonferroni correction were conducted in SPSS (IBMSPSS Statistics for Windows, version 25, SPSS Inc., Chicago, Ill., USA) with cue condition (4 levels: Valid-Valid, Valid-Invalid, Invalid-Valid, Invalid-Invalid) and target location on the vertical axis (2 levels: top, bottom) as within-participant factors, unless otherwise stated. When the ANOVA revealed a significant main effect or interaction, we computed the Partial eta squared 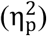 to estimate the effect size and ran Bonferroni tests for post-hoc comparisons. Correlation analyses were conducted with Pearson correlation tests. For all tests, the statistical significance was designated at *p*<0.05.

To test the statistical contrast in express response initiation time between the detrended-integrated signal and ROC analyses, we used single-subject statistical analysis (Supplementary Materials and Methods). For the detrended-integrated signal analysis, we also used this statistical approach to test the contrast in express response initiation time between the four different cue conditions at the single-subject level (Supplementary Materials and Methods; see 16 and 17 for further details).

## RESULTS

### Experiment 1

#### Task correctness, reaction time and kinematics

The cue validity had a significant effect on the prevalence of correct target-directed reaches (*F*_*3,14*_=23.3, *p*<0.001, 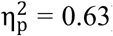). For both the top and bottom targets, the proportion of correct reaches was significantly higher when the target location was validly than invalidly cued (Table 1). Further, significantly fewer correct trials were observed in the Valid-Invalid than Invalid-Invalid cue conditions (Table 1). This indicates that providing invalid cues promoted incorrect reaching directions.

**Table 1:** Percentage of correct target-directed reaches toward top and bottom targets in the two experiments

|  |  | Cue conditions |  |  |  |
| --- | --- | --- | --- | --- | --- |
|  |  | Valid-Valid | Valid-Invalid | Invalid-Valid | Invalid-Invalid |
| **Experiment one** | Top target (%) | 98.8±1<sup>*,**,***</sup> | 76.1±16.6<sup>++</sup> | 82.9±10.9 | 86.6±9.8 |
|  | Bottom target (%) | 98.3±2.6<sup>*,**,***</sup> | 76.4±15<sup>++</sup> | 84.3±11.7 | 86.8±9.5 |
| **Experiment two** | Top target (%) | 96.3±3.5<sup>**,***</sup> | 95.9±4.4<sup>+,++</sup> | 89.6±6.4 | 86.6±9.2 |
|  | Bottom target (%) | 95.1±3.8<sup>**,***</sup> | 95.4±4.4<sup>+,++</sup> | 86.6±9.4 | 86.6±8.5 |

Statistically significant difference between the Valid-Valid cue condition and: \*, the Valid-Invalid cue condition; \*\*, the Invalid-Valid cue condition; \*\*\*, the Invalid-Invalid cue condition. Statistically significant difference between the Valid-Invalid cue condition and: +, the Invalid-Valid cue condition; ++, the Invalid-Invalid cue condition. Data reported as mean ± standard deviation.

The first line of panels in figure 4 shows exemplar correct reaching trajectories from a single participant toward the top-right target. For this person, the RT was ∼200ms when the target was validly cued, ∼240ms when only the right\left target location was validly cued and

**Figure 4:**
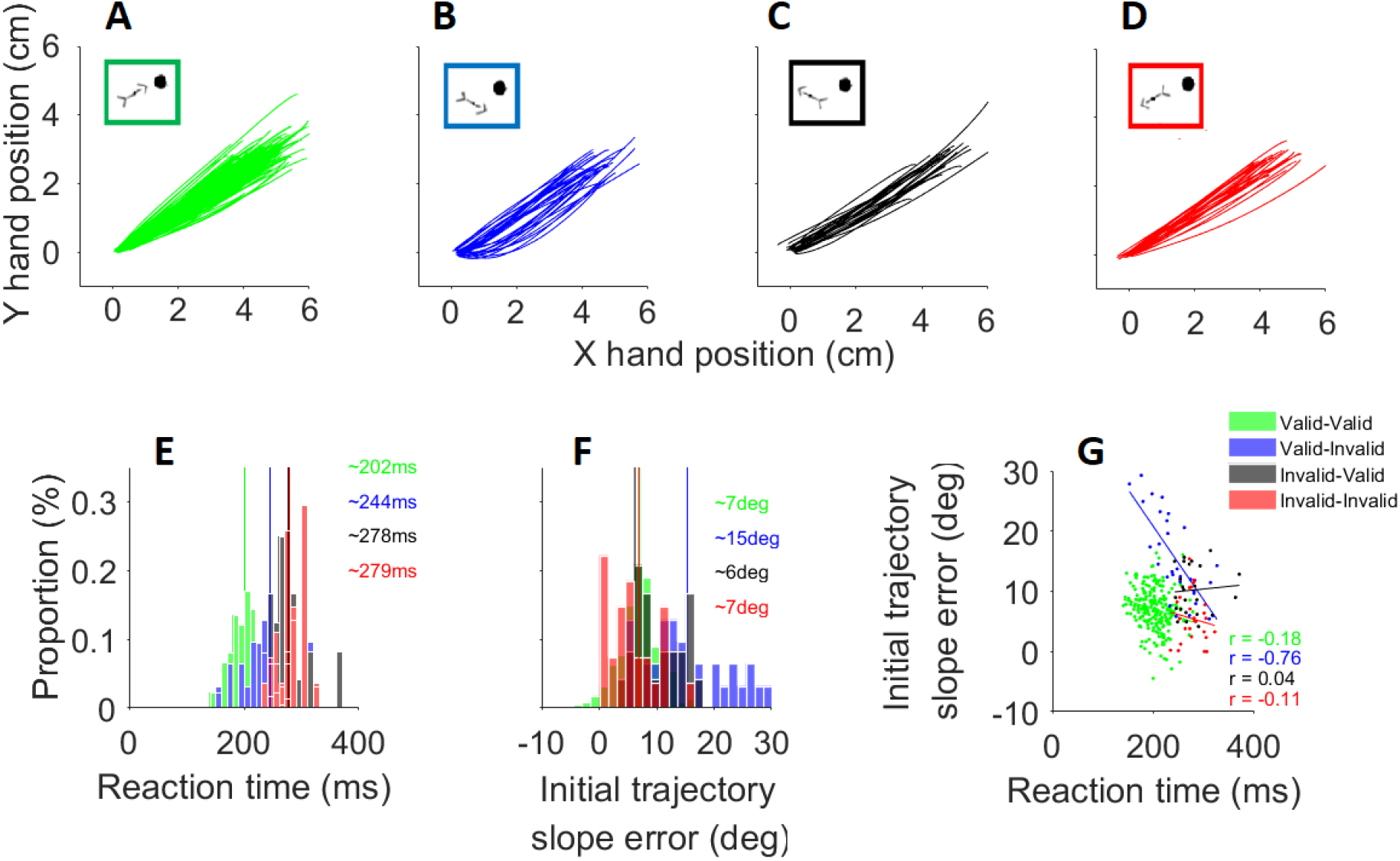
Correct target-directed reaching trials toward the top-right target of an exemplar subject from the first experiment. A, B, C and D panels show reaching trajectories for the Valid-Valid (green traces), Valid-Invalid (blue traces), Invalid-Valid (black traces) and Invalid-Invalid (red traces) cue conditions. For each cue condition, one variation of the cue orientation relative to the target location is shown in the framed squares. I, reaction time distribution for each of the four different cue conditions. E, distribution of the reaction time for each of the four different cue conditions. F, distribution of the initial trajectory angle errors for each of the four different cue conditions. G, relationship between the reaction time and the initial trajectory angle error (*r =* Pearson correlation coefficient).

∼280ms whenever the right\left cue orientation was invalid (Figure 4E). The initial trajectory angle errors were larger in the Valid-Invalid cue condition (∼15°) than in the other cue conditions (∼7°; Figure 4F). In the example used in figure 4B, this would indicate that the initial reach direction was biased more from the target direction (i.e. top-right) toward the cued direction (i.e. bottom-right) than was observed for the other cue conditions. Moreover, a strong negative correlation (*r = -*0.76; Figure 4G) between the initial trajectory angle error and RT was observed only for the Valid-Invalid cue condition, which indicates that the shorter the movement onset time the greater the cue-induced bias on the initial movement trajectory.

For the entire group, the RT was significantly influenced by the cue-condition (*F*_*3,14*_=75.3, *p*<0.001, 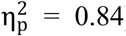) and the target location on the vertical axis (*F*_*1,14*_=28, *p*<0.001, 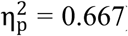), and by their interaction (*F*_*3,14*_=2.9, *p=*0.48, 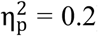). For both the top and bottom targets, the RT was significantly shorter when the target location was validly than invalidly cued (Figure 5A and D). The RT was also significantly shorter in the Valid-Invalid cue condition than the Invalid-Valid and Invalid-Invalid cue conditions (Figure 5A and D).

**Figure 5:**
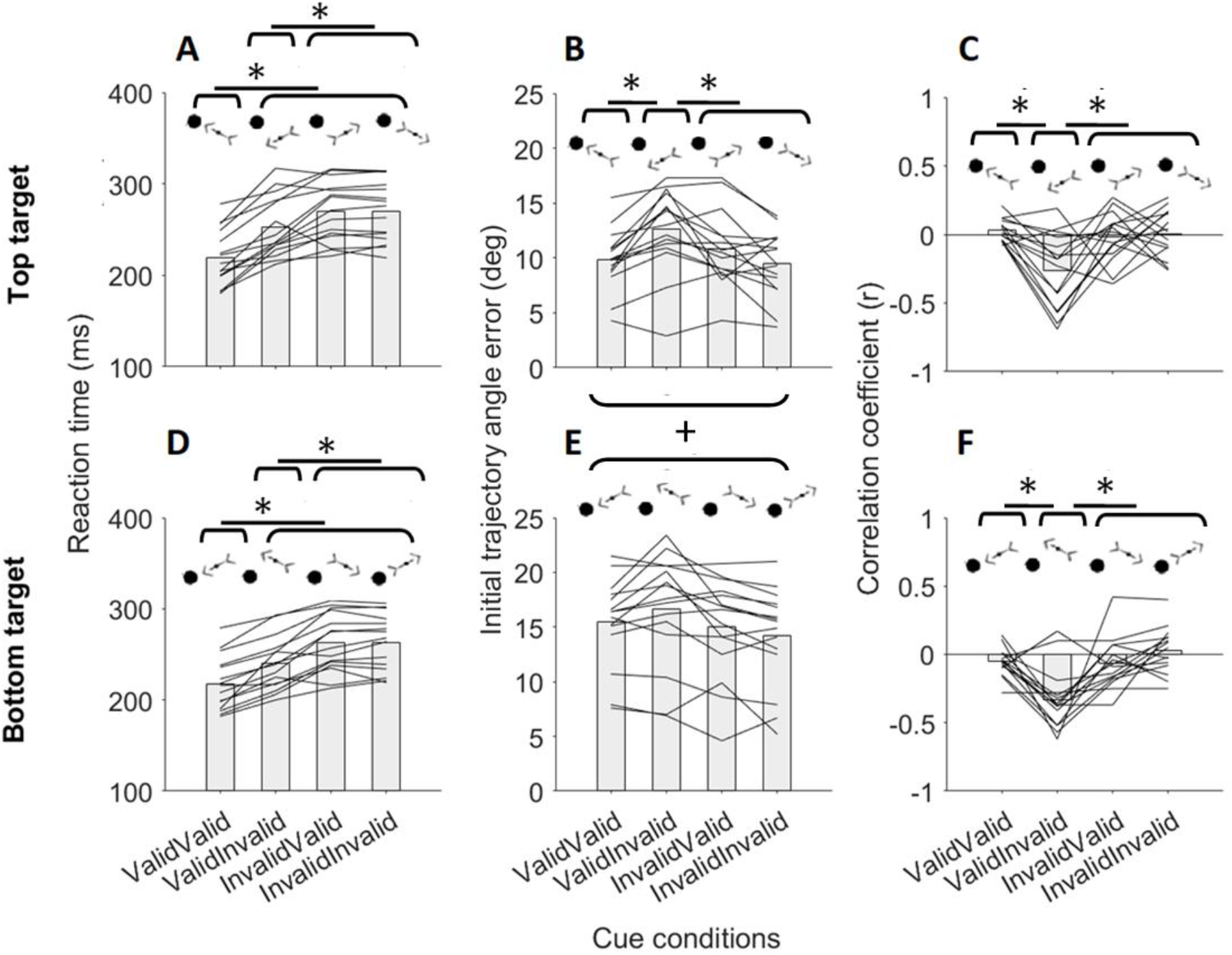
Metrics of the long-latency visuomotor behaviour of the first experiment. Latency of correct reaches in the four different cue conditions for the (A) top and (D) bottom targets. Angle error between the optimal and actual initial trajectory of the movement (see material and methods, and Figure 2 for details) in the four different cue conditions for the (B) top and (E) bottom targets. Correlation between the reaction time and the angle error of the initial trajectory (see material and methods, and Figure 2K) for the (C) top and (F) bottom targets. Each black line represents one participant and the bars represent the mean across subjects. Note that the exemplar cue conditions outlined above the bars represent only top (A, B, C) and bottom (D, E, F) left target conditions for clarity. *, statistically significant differences between the cue conditions; +, statistically significant differences between the target locations.

The initial trajectory angle error was significantly influenced by the cue-condition (*F*_*3,14*_=10.8, *p*<0.001, 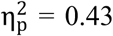) and by the target location on the vertical axis (*F*_*1,14*_=13.7, *p*=0.002, 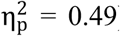), and by their interaction (*F*_*3,14*_=3.6, *p*=0.045, 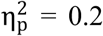). The positive trajectory error angles indicate that the initial movement was directed more horizontally than the optimal target-directed trajectory, especially for the bottom targets (significant differences between the top and bottom targets; Figure 5B and E). The initial trajectory error was larger in the Valid-Invalid cue condition than in the other cue conditions (Figure 5B and E), even though statistically significant pairwise contrasts were found only for the top target (Figure 5B). Further, the correlation coefficient between the initial trajectory error angle and the RT was significantly influenced by the cue-condition (*F*_*3,14*_=14.2, *p*<0.001, 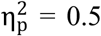), and was significantly more negative for the Valid-Invalid cue condition than the other cue conditions (Figure 5C and F).

In all, these data indicate that the valid right\left cues facilitated the RT. Validly cueing only the horizontal target location (i.e. Valid-Invalid cue conditions), however, also biased the initial movement toward the invalid top\bottom cued location. Critically, the initial trajectory angle error in the Valid-Invalid cue condition increased as participants reduced their RT, which justifies the larger prevalence of incorrect (i.e. non-target directed) responses for the Valid-Invalid than other invalid cue conditions.

#### Express muscle response

For the first experiment, twelve participants out of fifteen (80%) met the criteria for positive express response identification in each of the four cue conditions (see materials and methods) on the PMch muscle. By contrast, the conditions for positive express-response determination on the PD muscle among each different cue condition were not met by any participant. The discrepancy between the PMch and PD muscles is consistent with previous work (16, 17) and is probably due to the preload, which enhanced solely the activity of the PMch muscle. Given the low occurrence of express responses for the PD, only the PMch was considered for statistical analyses.

Figure 6 shows surface EMG recordings from the PM muscle of an exemplar express response producer from the first experiment. For this subject, the percentage of express response trials was higher when the right\left target location was validly (top target: Valid-Valid cue condition 69%, Valid-Invalid cue condition 65%; bottom target: Valid-Valid cue condition 75%, Valid-Invalid cue condition 72%) than invalidly cued (top target: Invalid-Valid cue condition 60%, Invalid-Invalid cue condition 44%; bottom-target: Invalid-Valid cue condition 64%, Invalid-Invalid cue condition 54%). For the top target, the muscle started encoding the target location at 90ms from its presentation when it was validly cued, at 95ms when only the right\left target location was validly cued (i.e. Valid-Invalid cue condition), and after 100ms when the target appeared opposite to the cued right\left visual hemi field (i.e. Invalid-Valid and Invalid-Invalid cue conditions). For the bottom target, the express response initiation time was ∼90ms when the right\left target location was validly cued (i.e. Valid-Valid and Valid-Invalid cue conditions), and >100ms when the right\left target location was invalidly cued (i.e. Invalid-Valid and Invalid-Invalid cue conditions). For both top and bottom targets, the express response initiation time in the Valid-Valid cue condition obtained via the detrended-integrated signal method was consistent with that resulting from the ROC analysis (Supplementary Figure 1A and B; Supplementary Table 1: experiment 1). For the top target, the express response magnitude was larger when the target location was validly cued (36µV) than in the other cue conditions (Valid-Invalid 27µV, Invalid-Valid 30µV, Invalid-Invalid 29µV). By contrast, for the target appearing close to the bottom of the monitor (i.e. bottom-target) the express response was 38µV in the Valid-Invalid cue condition, 32µV in the Valid-Valid cue condition and ∼25µV in the Invalid-Valid and Invalid-Invalid cue conditions.

**Figure 6:**
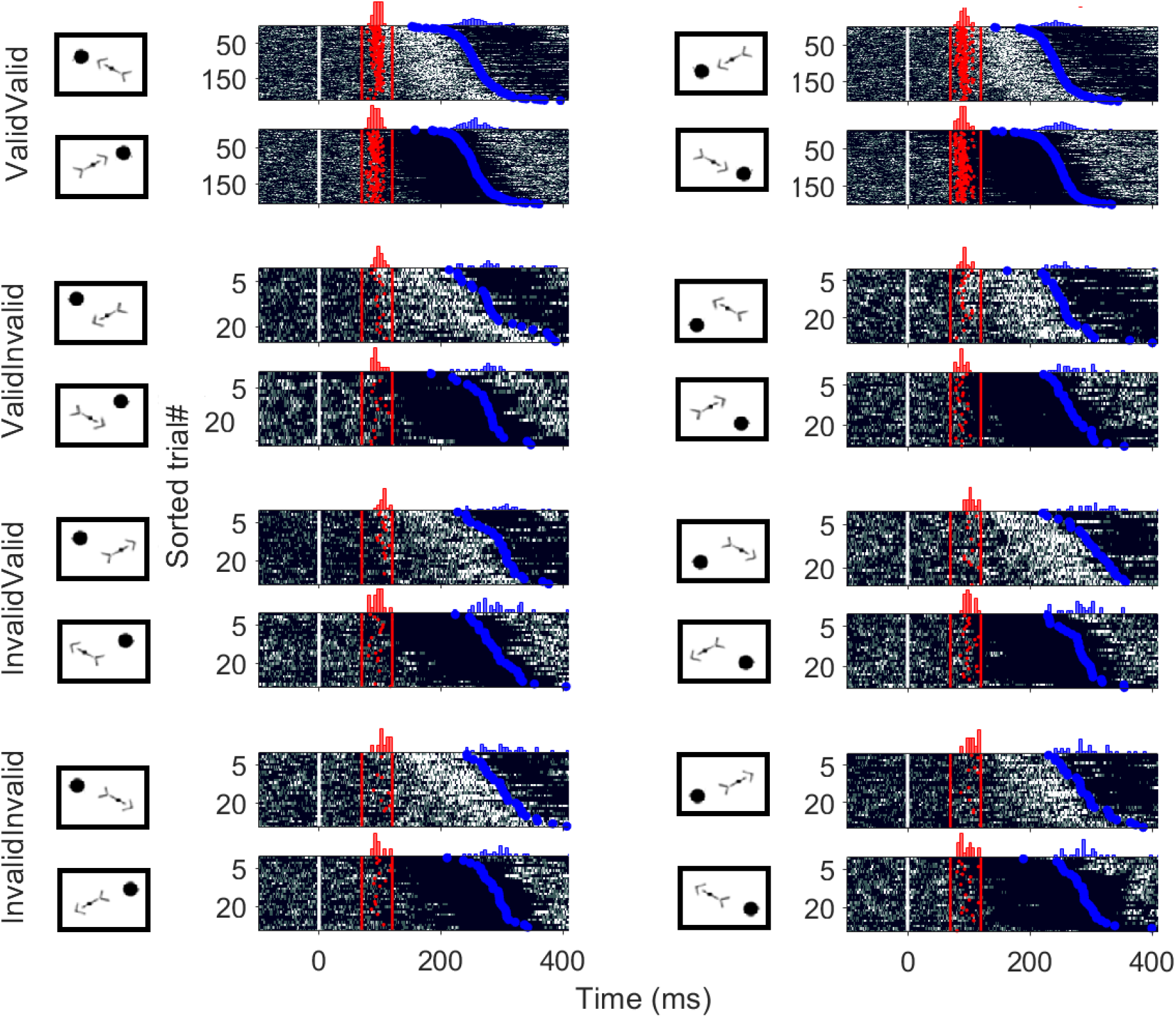
Surface EMG activity of the PM muscle during the leftward and rightward movements executed toward the top (first column of panels) and bottom (second column of panels) targets of an exemplar participant who completed the first experiment, and exhibited an express response in each of the four different cue conditions (see materials and methods). The first line of panels shows the condition in which the target location was validly cued (i.e. Valid-Valid cue condition). The second line of panels shows the condition in which only the right\left target location was validly cued (i.e. Valid-Invalid cue condition). The third line of panels shows the condition in which only the top\bottom target location was validly cued (i.e. Invalid-Valid cue condition). The fourth line of panels shows the condition in which the target was invalidly cued (i.e. Invalid-Invalid cue condition). For each cue condition, rasters of rectified EMG activity from individual trials are shown (same format as figures 3E-H; note that only express muscle response onset times are showed for clarity), as are the distributions of the muscle response initiation time (red histograms) and reaction time (blue histograms).

For the entire group, the prevalence of trials with an express muscle response was significantly influenced by the cue-condition (*F*_*3,11*_=16.7, *p*<0.001, 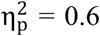) and the target-location on the vertical axis (*F*_*1,11*_=20.5, *p*=0.001, 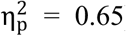), and by their interaction (*F*_*3,11*_=6.5, *p*=0.001, 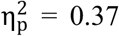). For the top target, the SLR prevalence was significantly higher in the Valid-Valid cue condition than the other cue conditions, and significantly higher in the Valid-Invalid than Invalid-Invalid cue conditions (Figure 7A). For the bottom target, the express response prevalence was significantly higher when the right\left target location was validly (Valid-Valid and Valid-Invalid cue conditions) than invalidly (Invalid-Valid and Invalid-Invalid cue conditions) cued (Figure 7D). The express response prevalence was also significantly higher for the bottom target than the top target, but only in the Valid-Invalid cue condition (Figure 7A and D).

**Figure 7:**
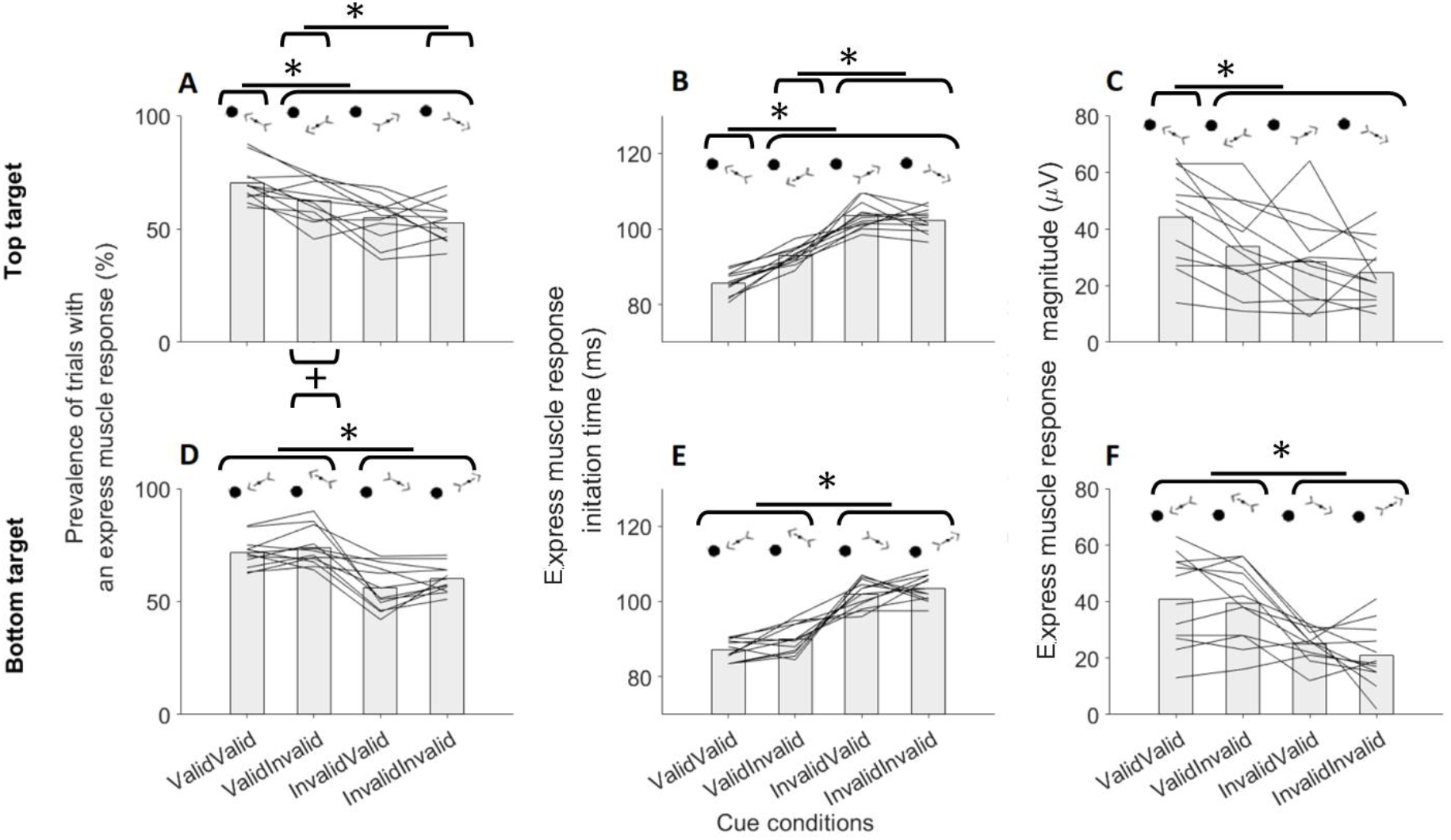
Express response metrics for the first experiment. Percentage of trials exhibiting an express muscle response (i.e. muscle response starting within 70-120ms from the visual stimulus presentation; see materials and methods) in the four different cue conditions for the (A) top and (D) bottom targets. Latency of the earliest stimulus-driven muscle response in the four different cue conditions for the (B) top and (E) bottom targets. Amplitude of the express visuomotor muscle response in the four different cue conditions for the (C) top and (F) bottom targets. Each black line represents one participant and the bars represent the mean across subjects. Note that the exemplar cue conditions outlined above the bars represent only top (A, B, C) and bottom (D, E, F) left target conditions for clarity. *, statistically significant differences between the cue conditions.

The differences in express response initiation time between the cue conditions were consistent among the twelve subjects who produced an express visuomotor response and resulted in statistically significant contrasts at the single-subject level (Supplementary Figure 2A and B; Supplementary Table 2: experiment 1). The ANOVA showed a significant cue-condition main effect (*F*_*3,11*_=103.1, *p*<0.001, 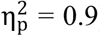), and a significant interaction between the cue condition and target location on the vertical meridian (*F*_*3,11*_=3.7, *p*=0.03, 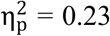). For both the top and bottom targets, the express response initiation time was significantly earlier when the right\left target location was validly (Valid-Valid and Valid-Invalid cue conditions) than invalidly cued (Invalid-Valid and Invalid-Invalid cue conditions; Figure 7B and E). The express response initiation time was also significantly shorter in the Valid-Valid cue condition than the Valid-Invalid cue condition, but only for the top target (Figure 7B). The express response magnitude was significantly influenced by the cue condition (*F*_*3,11*_=15.8, *p*<0.001, 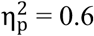).

For the top target, the express response size was significantly larger when the target was validly cued (Valid-Valid cue condition) than in the other cue conditions (Figure 7C). The express muscle response also tended to be larger in the Valid-Invalid than the Invalid-Valid and Invalid-Invalid cue conditions (Figure 7C), but these contrasts were not statistically significant. For the bottom target, the express response was significantly larger when the target appeared congruently (Valid-Valid and Valid-Invalid cue conditions) than incongruently (Invalid-Valid and Invalid-Invalid cue conditions) with the right\left orientation of the cue (Figure 7F).

Express response modulations secondary to valid and invalid horizontal cues could reflect facilitation and inhibition of early visuospatial processing, respectively. Express muscle responses, however, were not consistently modulated by the cue validity. Indeed, when the target was appropriate for the right\left cued location, invalidly cueing the top\bottom target location did not systematically impair the express response (see the results for the Valid-Invalid cue condition of bottom targets; second line of panels in Figure 7). Note that reaching for the top and bottom targets required different mechanical contributions from the muscle of interest. Both the top-left and bottom-left targets required a shoulder flexion in the transverse plane involving both the clavicular and sternal heads of the pectoralis major muscle. The top-directed movement, however, also required a shoulder sagittal plane flexion for which the clavicular head of the pectoralis is primarily involved. By contrast, the bottom-directed reach required a shoulder extension in the sagittal plane that mainly involves the sternal pectoralis fibres. We therefore tested if the cue-induced modulation of express responses was sensitive to the cued reaching direction, rather than the cue validity.

We ran a repeated measures ANOVA analysis on the express response magnitude to the left targets only in the Valid-Valid and Valid-invalid cue conditions. The ANOVA was conducted with Bonferroni correction, and with cue-orientation (2 levels: top orientation, bottom orientation) and target-cue compatibility (2 levels: compatible target, incompatible target) as within-participant factors. We found statistically significant main effects for both the cue-orientation (*F*_*1,11*_=9.9, *p*=0.009, 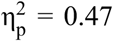) and target-cue compatibility (*F*_*1,11*_=10.6, *p*=0.008, 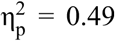). For both compatible and incompatible target-cue conditions, express muscle responses were significantly larger for the top than bottom cue orientation (Figure 8). The fact that the biomechanical action of the PMch aligns with the top-left target to a greater extent than the bottom-left target suggests that overt cue-driven expectation primes circuits that are sufficiently along the sensory to motor continuum to reflect biomechanical details of expected movement. For both cue orientations, the express response was significantly larger when the target location was compatible than incompatible with the cue orientation (Figure 8). This suggests that matching prior motor signals with spatially compatible visual inputs facilitated the generation of larger express visuomotor muscle responses.

**Figure 8:**
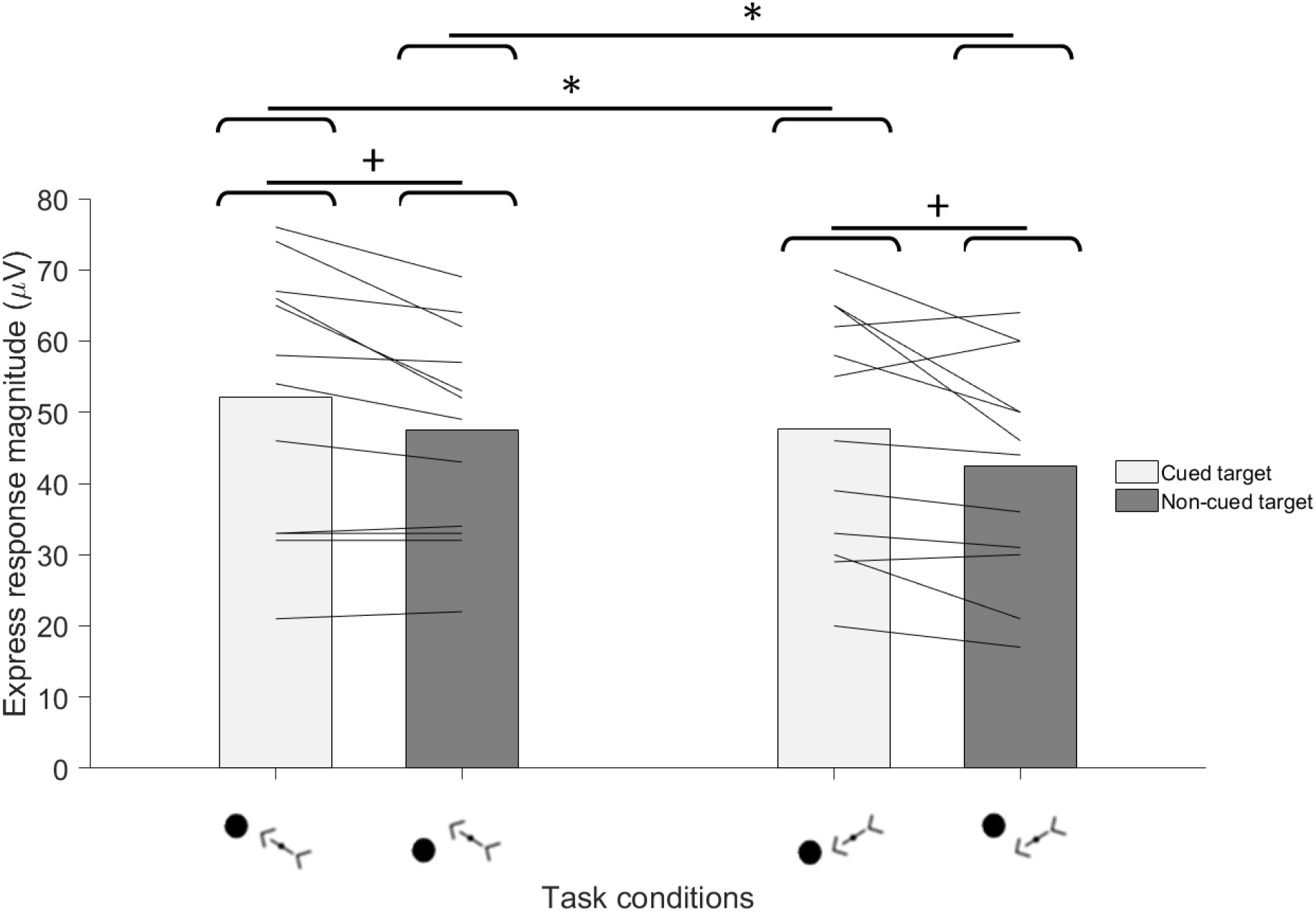
Dependency of the magnitude of express muscle response to the left targets on the cue orientation and its compatibility with the target location. Each black line represents one of the twelve subjects of the first experiment who exhibited an express response and the bars represent the mean across subjects. *, statistically significant differences between the cue orientation conditions; +, statistically significant differences between the cue-target compatibility conditions.

### Experiment 2

#### Task correctness and reaction time

The proportion of correct reaches was significantly influenced by the cue (cue-condition main effect: *F*_*3,15*_=20.02, *p*<0.001, 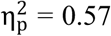). For both the top and bottom targets, the proportion of correct reaches was significantly higher when the reaching direction was validly cued than invalidly cued (Table 1). By contrast, the proportion of correct responses was not influenced by the top\bottom cue validity. These results suggest that the initial movement direction was biased solely by the right\left cue orientation.

The RT was significantly influenced by the cue validity (cue-condition main effect: *F*_*3,15*_=36.6, *p*<0.001, 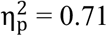). For both the top and bottom targets, the RT was significantly shorter when the reaching direction was validly (Valid-Valid and Valid-Invalid cue conditions) than invalidly (Invalid-Valid and Invalid-Invalid cue conditions) cued (Figure 9). This indicates that the participants used the right\left cue orientation to improve their performance, regardless of the top\bottom cue validity.

**Figure 9:**
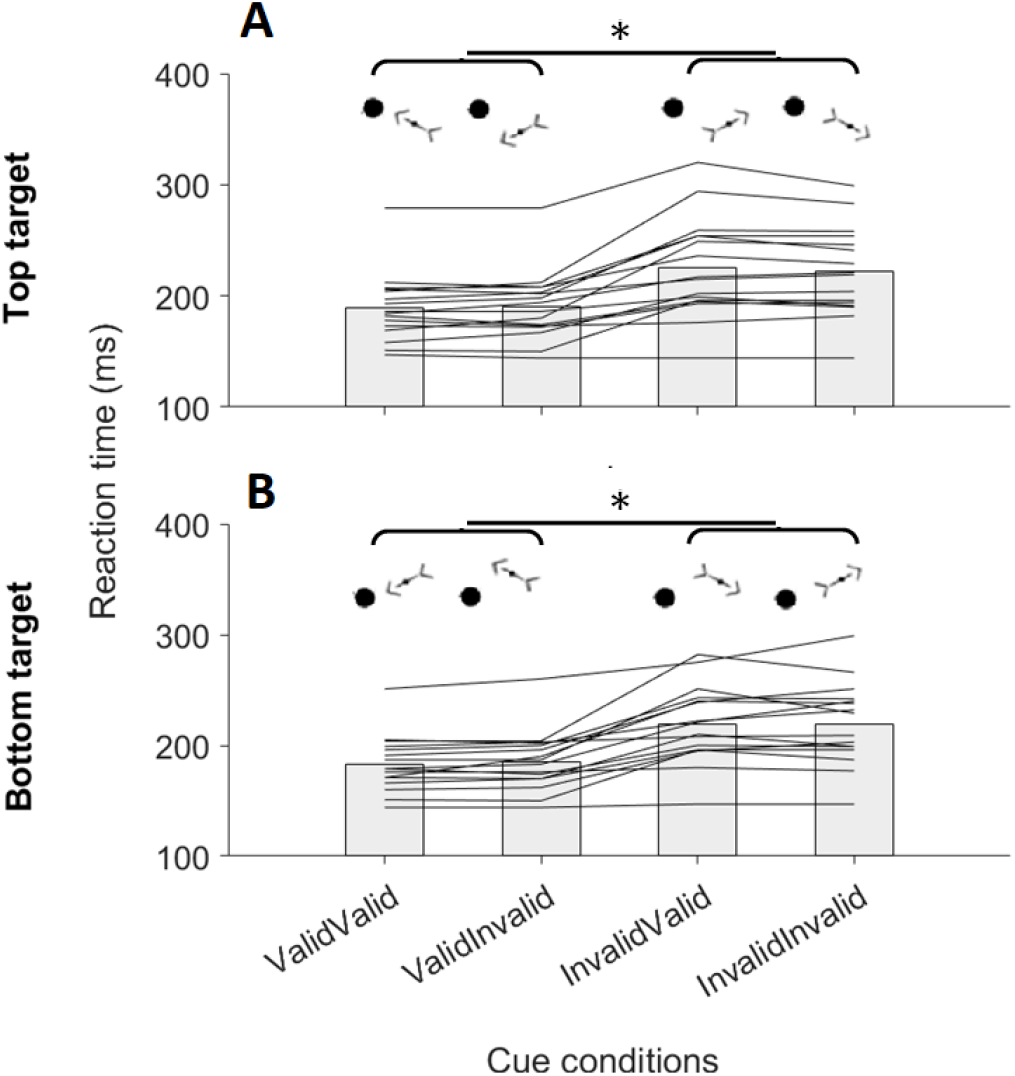
Reaction time of correct reaches in the four different cue conditions (see Figure 1A) for the (A) top and (B) bottom targets of the second experiment. Each black line represents one participant and the bars represent the mean across subjects. Note that the exemplar cue conditions outlined above the bars represent only top (A) and bottom (B) left target conditions for clarity. *, statistically significant differences between the cue conditions.

#### Express muscle response

Eleven out of the sixteen participants (i.e. ∼69%) in the second experiment exhibited an express visuomotor response on the PMch muscle on each of the four cue conditions. Again, no participant consistently exhibited a PD muscle express response across the different cue conditions and, thereby, only the express responses recorded on the PMch muscle were considered for statistical analysis.

Figure 10 shows the EMG recordings from one of the eleven subjects who showed an express response on the PMch muscle for each cue condition of the first experiment. The percentage of trials with an express muscle response was larger when the reaching direction was validly (Valid-Valid cue condition: top target 89%, bottom target 87%; Valid-Invalid cue condition: top target 84%, bottom target: 91%) than invalidly cued (Invalid-Valid cue condition: top target 36%, bottom target 57%; Invalid-Invalid cue condition: top target 52%, bottom target 48%). For both the top and bottom targets, the earliest target-driven muscle response was ∼93ms when the reaching direction was validly cued (i.e. Valid-Valid and Valid-Invalid cue conditions), and at ∼106ms when it was invalidly cued (i.e. Invalid-Valid and Invalid-Invalid cue conditions; see materials and methods).

**Figure 10:**
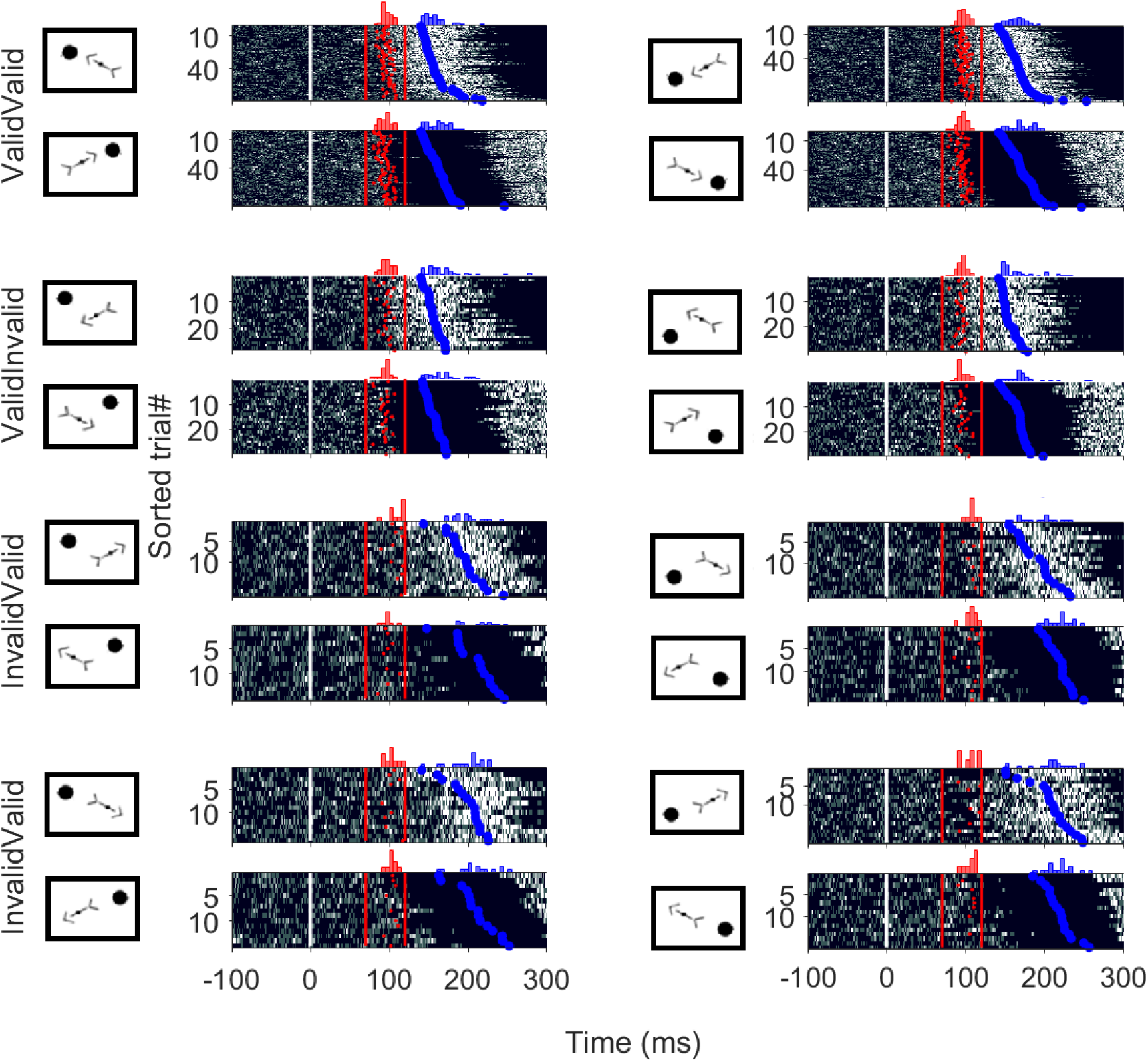
Surface EMG activity of the PM muscle during the leftward and rightward movements executed toward the top and bottom targets of an exemplar participant who completed the second experiment and exhibited an express response in each of the four different cue conditions (same format as figure 5).

Similar results were found among the eleven positive SLR subjects of the second experiment (Supplementary Figure 2C and D and Table 2: experiment 2). Again, the express response initiation times in the Valid-Valid cue condition were consistent among the detrended-integrated and ROC methods (Supplementary Figure 1C and D; Supplementary Table 1: experiment 2). The express response magnitude was larger when the reaching direction was validly (Valid-Valid cue condition: top target 108µV, bottom target: 127µV; Valid-Invalid cue condition: top target 114µV, bottom target: 114µV) than invalidly cued (Invalid-Valid cue condition: top target: 70µV, bottom target: 58µV; Invalid-Invalid cue condition: top target 51µV, bottom target 77µV).

For the entire group, the prevalence of express responses was significantly influenced by the cue conditions (*F*_*3,10*_=15.82, *p*<0.001, 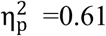). For the top target, the express response prevalence was significantly higher when the reaching direction was validly than invalidly cued (Figure 11A). Similar post-hoc results were observed for the bottom target, except that the contrast between the Valid-Invalid and Invalid-Valid cue conditions was not significant (*p*=0.09) (Figure 11D). The express response initiation time was significantly influenced by the cue condition (*F*_*3,10*_=159.79, *p*<0.001, 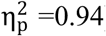). For both top and bottom targets, the express response initiation time was significantly earlier when the reaching direction was validly than invalidly cued, irrespective of the top\bottom cue validity (Figure 11B and E). A significant cue-condition main effect was also found for the express response magnitude (*F*_*3,10*_=11.13, *p*<0.001, 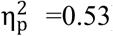). For the top target, the express response magnitude was significantly larger when the reaching direction was validly than invalidly cued (Figure 11C). For the bottom target, the express response magnitude was significantly larger in the Valid-Valid and Valid-Invalid cue conditions than the Invalid-Valid cue conditions (Figure 11F). These findings indicate that validly cueing the reaching direction positively modulated the express visuomotor behaviour, regardless of the prior information about the probable target location in the vertical axis.

**Figure 11:**
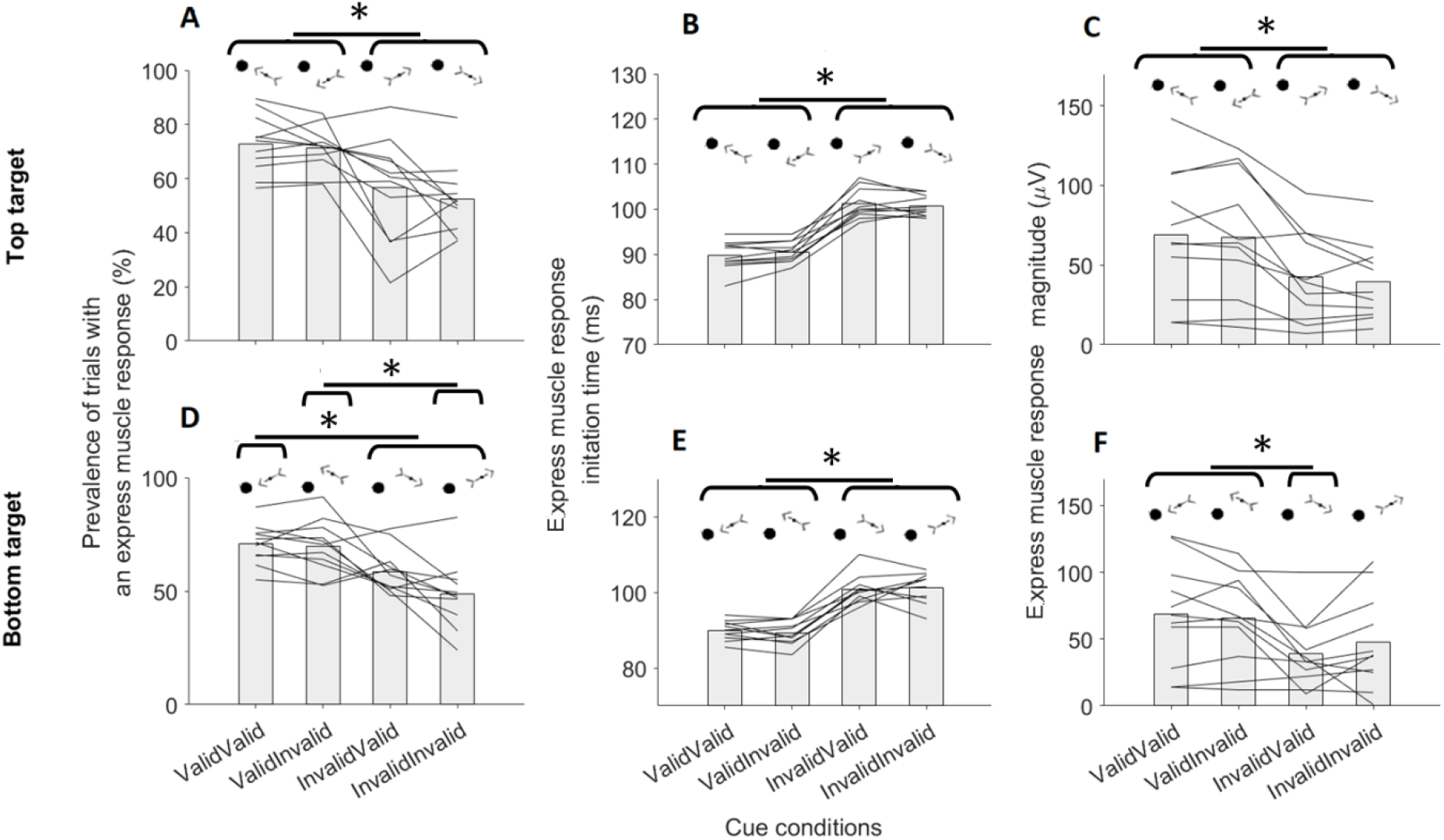
Prevalence, latency and magnitude of the express muscle response for the second experiment (same format as figure 6). *, statistically significant differences between the cue conditions; +, statistically significant differences between the target locations.

## DISCUSSION

Across the two experiments, we characterized cue-induced modulations of express arm muscle responses and long-latency kinematic parameters during a target-directed reaching task. Akin to express saccades (20, 21), express arm muscle responses result from rapid sensorimotor transformations of visual inputs into motor outputs. Further, express muscle responses appear to lack the flexibility to implement task rules as they consistently encode the physical location of visual targets within ∼100ms from their presentation even in anti-reach (7) or no-reach (10) tasks. This implies a control network with properties similar to that for express saccades, which involves the midbrain superior colliculus and the brainstem reticular formation (30, 31). Considering that the participants had to interpret symbolic meanings of cue orientations, our results suggest that even the earliest neural computations required to transform visual inputs into motor outputs are sensitive to modulation reflecting top-down expectations. Notably, the time for sensory-to-motor transformation of visual inputs along the express pathway was ∼15ms shorter with valid than invalid cues.

### Cue-driven visuospatial facilitation

In the first experiment in which the participants could execute four distinct target-directed reaches, validly cueing the target location facilitated express visuomotor behaviours relative to the invalid cue conditions. This could reflect an overt cue-induced mechanism for orienting attention (32) or an attentional bias driven by motor intention, a phenomenon known as the *intentional weighting mechanism* (33-35). In this circumstance, the sensory-to-motor transformation of visual inputs would be facilitated and inhibited for validly and invalidly cued targets, respectively. Notably, there are extensive cortico-tectal projections () that are well-placed to mediate the top-down delivery of cortical (e.g. cue-driven) signals to the superior colliculus (37, 38), and modulate the sensorimotor transformations operated by this midbrain structure. This is consistent with evidence of reduced collicular activity (39) and express saccade prevalence (40) after cryogenic inactivation of the frontal eye field in monkeys. Earlier behavioural work also showed that express visuomotor behaviour is modulated as a function of temporal and spatial stimulus predictability (16, 17, 20, 41), and explicit cue-driven instructions (42). Overtly cueing the target location might facilitate visuaospatial processing of stimuli at the cued locations on the collicular visual map such that the visual signals encoding expected targets could be more rapidly integrated within visuomotor circuits that project to the motor nuclei of the reticular formation (see 19 for review).

Although the target appeared at each of the three invalidly cued locations with equal probability in the first experiment, express muscle responses were facilitated for the Valid-Invalid cue condition relative to the other invalid cue conditions. This might reflect a broadly distributed facilitation of the colliculus encoding the cued right\left visual hemi field, possibly by reducing excitatory drive to the foveal fixation zone that has non-specific inhibitory projections to all extrafoveal parts of that colliculus (see for review 19, 36). Basso and Wurtz (43), however, showed that validly cueing one of eight possible targets (45° between the targets; ∼10dva of eccentricity from fixation) facilitated only the collicular neurons encoding the cued locus. Some facilitation of collicular visual map restricted at the cued locus should have occurred also in our task because we used an even larger (∼60°) top-bottom target gap. Furthermore, the cue-induced modulations of express muscle response were not symmetrical around the horizontal axis. Specifically, providing only right\left valid information (i.e. Valid-Invalid cue condition) impaired express muscle responses to the top targets, but not bottom targets, relative to valid cue conditions (i.e. Valid-Valid cue condition). This asymmetry argues against a generalised cue-driven left/right hemifield enhancement of attention.

### Cue-driven motor facilitation

Larger express muscle responses resulted from validly cueing the location of top-left than bottom-left targets (see first and third bars in Figure 8) of the first experiment. Although this could reflect functional discontinuity of the superior colliculus across the horizontal axis (44), our results for invalid top\bottom cue conditions suggest a different mechanism. Specifically, larger express muscle responses were detected by invalidly cueing the bottom-left (i.e. cue oriented top-left) than top-left (i.e. cue oriented bottom-left) target (see second and fourth bars in Figure 8). Notably, this is in line with the mechanical output required to the PMch to bring the hand at top and bottom target locations (see results section).

The pectoralis major muscle has multiple zones of innervations that are confined medially within the cranio-caudal muscle length (45). This subserves discrete recruitment of different muscle portions to accomplish the required movement depending on the posture and the upper limb osteo-kinematics (46-48). Critically, the compatibility between express muscle responses and cued reaching directions suggests that the cue-induced modulations of express muscle responses were inclusive of prior cue-driven motor signals, rather than solely sensitive to facilitation of rapid visuospatial target processing.

The cue-induced motor preparation mechanism was further tested in the second experiment in which a single rightward or leftward reach was required. Here, validly cueing the right\left component of the target location facilitated express muscle response to both the top and bottom targets to a similar degree, irrespective of the top\bottom cue validity. Again, this is not consistent with evidence of collicular functional asymmetry across the horizontal axis (44), but rather appears to reflect a mechanism by which a prepared (i.e. cued) motor response can be rapidly released by any stimulus appearing compatibly with the expected reaching direction. By contrast, express muscle responses were likely impaired in invalid right\left cue conditions because of additional neural computations required to invert the agonist\antagonist muscle contribution to the prepared motor plans within the putative subcortical express pathway. Notably, our results are consistent with previous work by Gu et al. (8) in which straight target-directed reaches were encumbered by virtual obstacles that were presented one second before the target, thus prompting pre-target preparation of curvilinear reaches. The magnitude of express muscle response was similar for targets requiring curved reaching trajectories (e.g. right-then-top) and for those presented in line with the initial phase of the planned curved movement (e.g. right), whereas incongruent targets led to weaker express arm muscle responses.

Note that a cue-induced mechanism for motor preparation is also consistent with the cue-induced modulations of RT and kinematics. In the first experiment, providing only valid horizontal cues (i.e. Valid-Invalid cue conditions) biased the initial movement trajectory toward the invalid top\bottom cued direction. This led to significant reduction in task accuracy and increase in initial trajectory angle error for the Valid-Invalid cue condition relative to the other cue conditions. Furthermore, the negative correlation observed between the trajectory angle error and the RT in the Valid-Invalid cue condition suggests a progressive shift from cue-driven to target-driven motor plans. This is consistent with evidence of time-based inhibition of non-target directed motor responses, such as those encoding the location of distractor stimuli. Specifically, the occurrence of distractor-directed motor responses depends on the time available to suppress the distractor-driven motor signals (see 49 for review). The large initial trajectory angle errors in the Valid-Invalid cue conditions might, therefore, reflect under-inhibited motor plans to reach the invalid top\bottom cued location due to short delay between target presentation and movement initiation. This may reflect a situation in which a response is initiated as soon as the compatibility between pre-stimulus and post-stimulus motor plans is good enough to achieve the task goal, consistent with a trade-off between rapid versus efficient (shorter trajectories → lower energetic costs) responses. By contrast, the long RTs recorded in the invalid right\left cue conditions could reflect additional neural computations to transform the visual stimuli into unprepared motor signals and inhibit the cue-driven motor plans to avoid wrong movements. The results of the second experiment are also consistent with a cue-induced mechanism for motor preparation, since reaches were facilitated (inhibited) for any visual input compatible (incompatible) with the cued reach direction, regardless of cue-driven sensory expectations.

Classical work involving single unit brain recordings from monkeys indicated that multiple potential target directions are represented simultaneously in premotor cortices (50) and the superior colliculus (43), and that these representations compete for execution until the actual target location is known. More recent evidence from simultaneous recordings of multiple units in premotor cortex (51), however, suggests that the brain prepares only one motor plan than can be rapidly modified if necessary once the actual target location is known (51, 52). Notably, the cortical areas involved in pre-target motor planning are mutually interconnected with those encoding quantity, probability and value information (49, 53) that can facilitate the decision of which movement to prepare. Cueing the probable target location may trigger a process whereby the brain prepares a motor plan consistent with prior cue-driven information, and then integrates the cue-driven and target-driven motor signals to produce the final motor output (54, 55). The cortical signals affording cue-induced motor expectations might set the collicular visuomotor state (43) via the cortico-tectal projections (36-38), thus modulating the neural computations to integrate cue-driven and target-driven motor plans operated by this midbrain structure. Downstream from the superior colliculus, the brainstem reticular formation receives extensive descending projections from the cortical motor areas. In particular, Keizer and Kuypers (56, 57) showed direct cortico-reticular projection originating from primary motor and premotor cortices in cats and monkeys. More recent studies extended this knowledge by showing both ipsilateral and contralateral cortico-reticular projections from the primary motor cortex (58, 59), the supplementary motor area (58, 59, 60) and the premotor cortex (58, 59) to the motor nuclei of the reticular formation. These cortico-reticulo projections might subserve the top-down delivery of preparatory motor signals and modulate the reticular formation motor set prior to the stimulus presentation. Overall, our findings are consistent with existing ideas about the role of the reticular formation in the rapid release of prepared motor actions (61, 62) at the arrival of triggering signals (e.g. from the superior colliculus) that arise from the presentation of salient stimuli.

### Functional motor behaviour implications of express muscle responses

It is worth nothing that the earliest volitional visuomotor behaviour appears to be facilitated by express muscle responses. We and others previously showed that larger express muscle responses were associated with earlier mechanical responses in humans (5, 7, 16, 17). Furthermore, Gu and colleagues (7) recorded express (∼110ms) hand-force divergences encoding the target location whose amplitude correlated with that of express muscle responses. Critically, the initiation time of hand-force divergence is consistent with a ∼30ms electromechanical delay (63) from the earliest stimulus-driven muscle response. Therefore, even though express muscle responses may not by themselves produce enough muscle force to reach the threshold for RT detection, they will still alter the muscle mechanical state to facilitate the subsequent volitional train of action potentials from spinal motoneurons.

### Conclusions

The present work documents modulations of express visuomotor arm muscle responses due to explicit cue-driven information in humans. The modulations of express responses appear to reflect a cue-driven mechanism for motor preparation within the putative subcortical express pathway, potentially including the midbrain superior colliculus and brainstem reticular formation. This might subserve the rapid and robust release of prepared motor responses to predictable visual events. Overall, our data lend support to the idea that there are meaningful subcortical contributions to visually-guided arm functions in humans, and that putative subcortical express pathway is subject to cortical modulation that enhances behavioural flexibility.

## Acknowledgements

This work was supported by operating grants from the Australian Research Council (DP170101500) awarded to T.J. Carroll, B.D. Corneil, G.E. Loeb and G. Wallis.

