## Supplementary Materials for "Cue-driven motor planning facilitates express visuomotor responses in human arm muscles"

Samuele Contemori,[1] Gerald E. Loeb,[2] Brian D. Corneil,[3,4,5] Guy Wallis,[1] Timothy J. Carroll[1]

1. Centre for Sensorimotor Performance, School of Human Movement and Nutrition Sciences, The University of Queensland, Brisbane, Australia.
2. Department of Biomedical Engineering, University of Southern California, Los Angeles, California, USA.
3. Department of Physiology and Pharmacology, Western University, London, Ontario, Canada.
4. Department of Psychology, Western University, London, Ontario, Canada.
5. Robarts Research Institute, London, Ontario, Canada.

### Supplementary Materials and Methods

In this work, we investigated the cue-induced modulations of express visuomotor muscle response by testing four different cue conditions across two experiments (see materials and methods of the main manuscript for details).

To extrapolate the earliest stimulus-driven muscle response, we developed a trial-by-trial approach named the *detrended-integrated* signal method (see materials and methods of the main manuscript for details). In addition, we compared our the detrended-integrated signal method with a time-series *receiver operator characteristics* (ROC) analysis that we and others previously used to determine the muscle response onset time (Pruszynski et al., 2010; Gu et al., 2016; Gu et al., 2018; Gu et al., 2019; Kozak et al., 2019; Kozak et al. 2020; Kozak and Corneil 2021; Contemori et al. 2021a and 2021b; see materials and methods of the main manuscript for details). Note that this analysis method considers simultaneously all valid trials within each condition. Briefly, we computed the area under the ROC curve (AUC) on each data sample recorded between 100ms before to 300ms after the target onset time. The AUC indicates the probability that an ideal observer could discriminate the target position from the EMG trace, with 0 and 1 representing perfect discrimination and 0.5 representing chance discrimination. A candidate express stimulus-locked distribution of trials was defined if the AUC overcame the 0.65 threshold within 70-120ms after target presentation, and remained above that threshold level for at least 15ms (Contemori et al. 2021a and 2021b). To test if the express muscle response was consistently time-locked to the stimulus onset time, we ran the ROC analysis either on the fastest or slowest halves of trials that were obtained by doing a median split on the RT data. We then associated the fast and slow candidate initiation times with the average RT of fast and slow data sets, and we fitted a line to the data to test if the earliest muscle response co-varied with the RT (i.e. line slope <67.5°; for further details see Contemori et al. 2021a and 2021b). In this case, we ran the ROC analysis on all trials and used a two-pieces linear regression analysis to determine the point in time at which the ROC curve starts diverging from background toward the 0.65 discrimination threshold, which defined the initiation time of the express visuomotor response (for further details see Contemori et al. 2021a and 2021b).

Given that the ROC analysis has low reliability for small data samples (Cross et al. 2019), this analysis was ran only for the cue condition in which the largest number of trials was recorded (i.e. Valid-Valid cue condition; see materials and methods of the main manuscript for further details).

*Identifying statistically significant contrasts at the single-subject level*

To test the statistical contrast in express response initiation time between the detrended-integrated signal and ROC analyses, we used a single-subject statistical analysis (Contemori et al. 2021a and 2021b) using Matlab (version R2018b, TheMathWorks, Inc., Natick, Massachusetts, United States). Specifically, we generated one thousand randomly re-sampled (with replacement) trial sets by using a bootstrapping approach on the original data and ran the two methods on each bootstrapped data set. This allowed us to determine the express response initiation time distribution for each of the two methods. We then compared one randomly re-sampled set of values obtained with the ROC analysis with one randomly re-sampled set of values obtained with the detrended-integrated signal analysis. If the values for one method were larger or smaller than those for the other method in more than 95% (i.e. >950) of cases, we concluded that the difference between the two methods was statistically significant (i.e. $p< 0.05$; for further details see Contemori et al. 2021a and 2021b).

For the detrended-integrated signal analysis, we also ran the single-subject statistical analysis on the express response initiation time across the four different cue conditions of the two experiments.

**Supplementary Results: Experiment 1**

*Between-methods contrasts*

For the first experiment, we compared the two methods of analysis on each of the twelve positive express responses producers (see materials and methods of the main manuscript for details), thus including the exemplar subject's data shown in figure 6 of the main manuscript. For this subject, the detrended-integrated signal analysis on the original data set showed that the express muscle response onset time in the Valid-Valid cue condition was ~90ms both for the top and bottom targets. The result for the top target result was consistent with those observed by running the detrended-integrated signal and ROC analyses on 1000 bootstrapped data sets (Supplementary Figure 1A; Supplementary Table 1: experiment 1, subject 7). For the bottom target, the express response onset time detected with detrended-integrated signal on 1000 bootstrapped data sets was ~2ms earlier than that obtained with the ROC analysis, and this difference was statistically significant ($p<0.05$; Supplementary Figure 1B; Supplementary Table 1: experiment 1, subject 7). Note, however, that this statistically significant contrast was shown for completeness because it was found only for four out of twenty-four recordings (Supplementary Table 1: experiment 1).

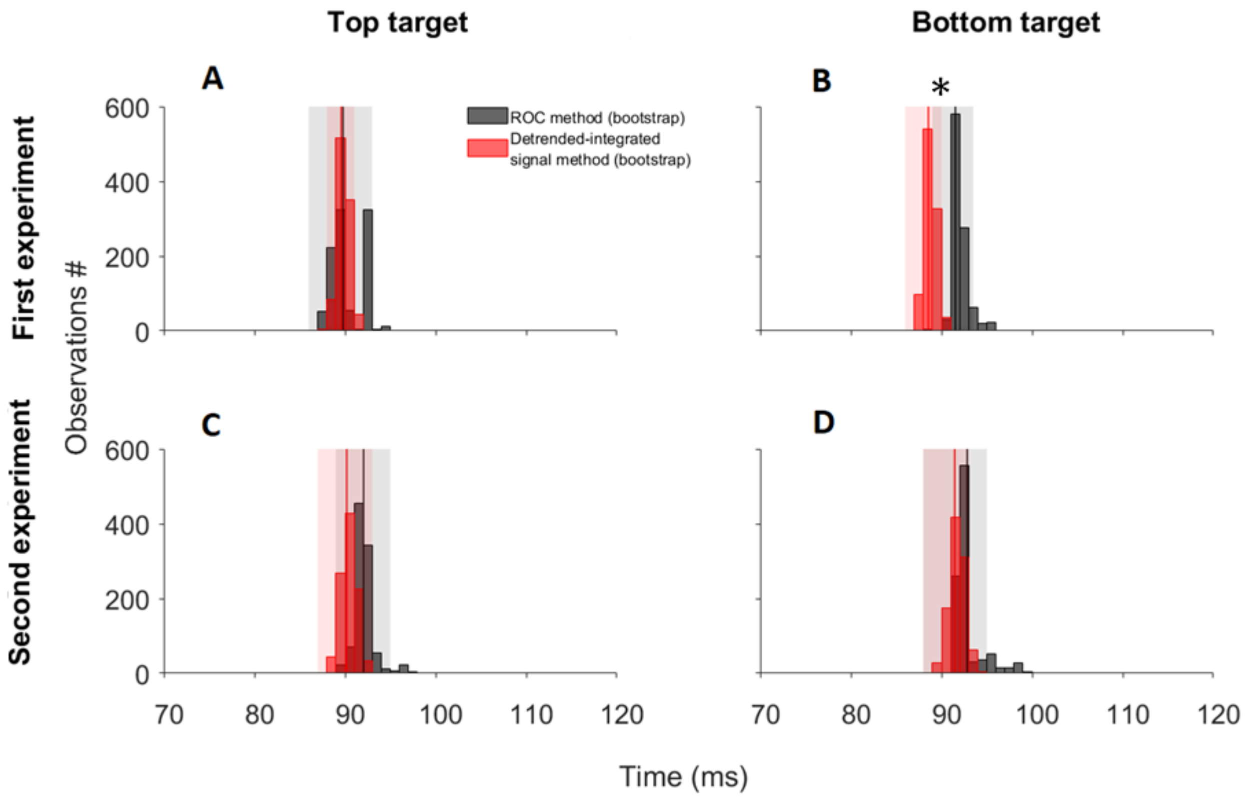


**Supplementary Figure 1:** Panels A and B show the distribution of express muscle response onset time of the exemplar subject presented in figure 6 of the main manuscript (Supplementary Table 1: experiment 1, subject 7). Panels C and D show the data of the exemplar subject presented in figure 10 of the main manuscript (Supplementary Table 1: experiment 2, subject 4). The panels show the pairwise comparisons between the detrended-integrated signal and ROC methods for both top and bottom targets. On each panel, the express response onset time distributions obtained by running the detrended-integrated signal and ROC analyses on 1000 bootstrapped data sets are shown, as are the 95% distribution confidence interval (patches) and the average distribution value (solid vertical lines). *, statistically significant differences between the methods.

**Supplementary Table 1**: Results of the single-subject statistical analysis on the muscle response onset times (in milliseconds) obtained by running the *detrended-integrated* signal analysis and ROC analysis on 1000 bootstrapped data sets (see materials and methods).

| **Target location** | Top target | | Bottom target | |
|---|---|---|---|---|
| **Analysis method** | Detrended-integrated signal | ROC | Detrended-integrated signal | ROC |
| Experiment 1 | | | | |
| Subject | | | | |
| 1 | 81 (80, 82) | 82 (78, 85) | 83 (82, 85) | 85 (82, 88) |
| 2 | 82 (81, 83) | 79 (74, 84) | 84 (83, 85) | 84 (80, 88) |
| 3 | 90 (88, 92)* | 95 (91, 99) | 90 (89, 92) | 87 (81, 92) |
| 4 | 88 (86, 89) | 86 (83, 90) | 90 (88, 91) | 85 (78, 92) |
| 5 | 86 (84, 88) | 88 (84, 92) | 88 (87, 89) | 89 (86, 91) |
| 6 | 82 (81, 83) | 85 (76, 94) | 84 (82, 85) | 91 (81, 100) |
| 7 | 90 (88, 91) | 90 (86, 93) | 88 (86, 90)* | 91 (89, 92) |
| 8 | 86 (84, 87) | 88 (82, 93) | 86 (84, 87) | 81 (76, 87) |
| 9 | 86 (84, 87) | 81 (71, 91) | 90 (88, 91) | 89 (78, 100) |
| 10 | 88 (87, 90)* | 98 (95, 100) | 90 (89, 92) | 92 (88, 96) |
| 11 | 85 (84, 86) | 82 (75, 90) | 86 (85, 88) | 85 (79, 91) |
| 12 | 85 (83, 87) | 79 (70, 88) | 85 (84, 87)* | 78 (71, 86) |
| Experiment 2 | | | | |
| Subject | | | | |
| 1 | 88 (86, 90) | 86 (83, 89) | 90 (88, 92) | 89 (85, 94) |
| 2 | 94 (92, 96) | 91 (82, 99) | 94 (92, 96) | 91 (86, 96) |
| 3 | 90 (88, 92) | 93 (88, 98) | 90 (88, 92) | 92 (86, 98) |
| 4 | 90 (87, 93) | 92 (89, 94) | 91 (90, 93) | 92 (88, 95) |
| 5 | 89 (87, 91)* | 105 (92, 117) | 89 (87, 91)* | 101 (82, 120) |
| 6 | 87 (85, 89) | 88 (81, 96) | 87 (85, 89) | 89 (85, 93) |
| 7 | 91 (89, 94)* | 101 (97, 104) | 91 (89, 93)* | 100 (95, 105) |
| 8 | 90 (88, 92) | 91 (87, 94) | 89 (86, 91) | 91 (87, 96) |
| 9 | 88 (86, 90) | 92 (82, 102) | 88 (83, 87)* | 96 (89, 102) |
| 10 | 83 (81, 85) | 92 (78, 106) | 85 (83, 87) | 89 (81, 97) |
| 11 | 88 (86, 91) | 92 (84, 100) | 87 (84, 89) | 88 (80, 96) |

First experiment's subjects 1, 4, 2, 8, 5, 7, 12 and 13 correspond to second experiment's subjects 1-5, 7, 10 and 11. The data are reported as mean (95% confidence interval). *, statistically significant differences between the two methods.

#### *Between-cue conditions contrasts*

For the second experiment, we compared the express muscle response initiation time between the four different cue conditions for each of the twelve positive express responses producers (see materials and methods of the main manuscript for details), thus including the exemplar subject's data shown in figure 6 of the main manuscript. For this subject, the detrended-integrated signal analysis on the original data set showed that the express muscle response onset time for the top target was 90ms when it was validly cued, at 95ms when only the right\left target location was validly cued (i.e. Valid-Invalid cue condition), and after 100ms when the target appeared opposite to the cued right\left hemi visual field (i.e. Invalid-Valid and Invalid-Invalid cue conditions). For the bottom target, the express response initiation time was ~90ms when the right\left target location was validly cued (i.e. Valid-Valid and Valid-Invalid cue conditions), and >100ms when the right\left target location was invalidly cued (i.e. Invalid-Valid and Invalid-Invalid cue conditions). These results were consistent with those observed by running the detrended-integrated signal analysis on 1000 bootstrapped data sets (Supplementary Figure 2A and B; Supplementary Table 2: experiment 1, subject 7). For the top target, the single-subject analysis showed that the express muscle response onset time was significantly earlier ($p<0.05$) when the target was validly than invalidly cued. In addition, the express muscle response onset time was significantly earlier ($p<0.05$) when in the Valid-Invalid than the Invalid-Valid and Invalid-Invalid cue conditions (Supplementary Figure 2A and B; Supplementary Table 2: experiment 1, subject 7). For the bottom target, the single-subject analysis showed that the express muscle response onset time was significantly earlier ($p<0.05$) when the right\left target location was validly (i.e. Valid-Valid and Valid-Invalid cue conditions) than invalidly (i.e. Invalid-Valid and Invalid-Invalid cue conditions) cued (Supplementary Figure 2A and B; Supplementary Table 2: experiment 1, subject 7). The results of the exemplar subject were also consistent among the twelve positive express response producers of the second experiment (Table 2: experiment 1).

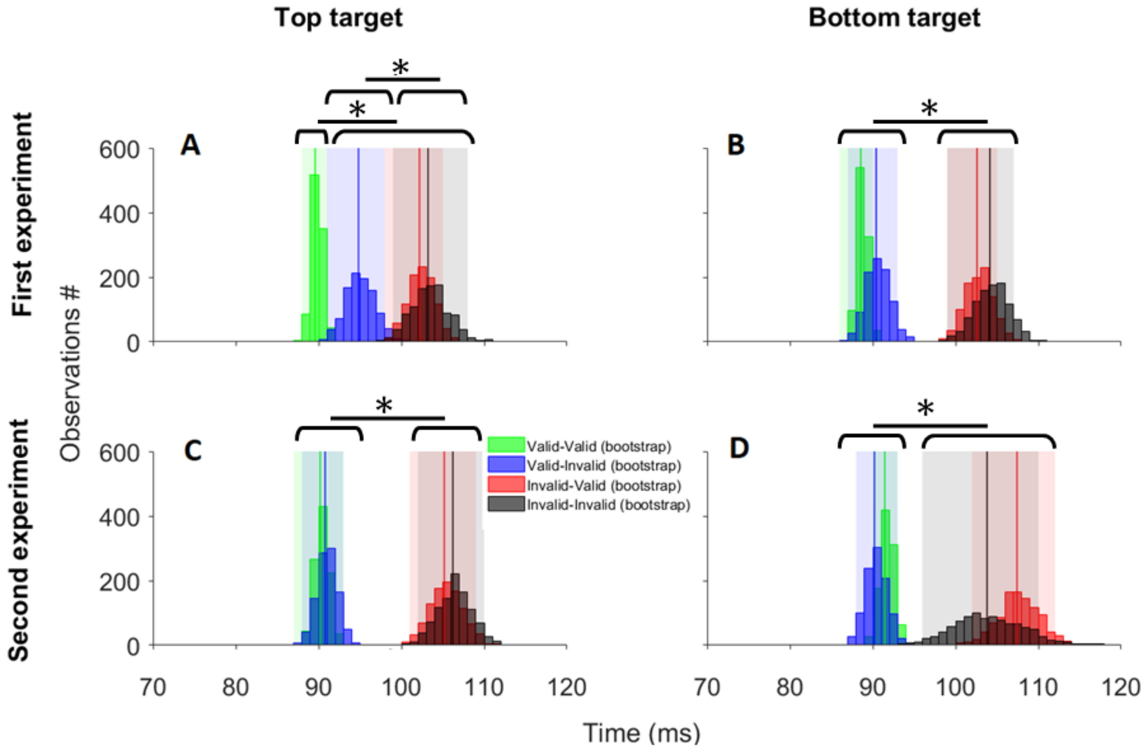


**Supplementary Figure 2:** Panels A and B show the distribution of express muscle response onset time of the exemplar subject presented in figure 6 of the main manuscript (Supplementary Table 2: experiment 1, subject 7). Panels C and D show the data of the exemplar subject presented in figure 10 of the main manuscript (Supplementary Table 1: experiment 2, subject 4). The panels show the comparisons between the four different cue conditions for both top and bottom targets (same format of Supplementary Figure 1). *, statistically significant differences between the cue conditions.

**Supplementary Table 2**: Results of the single-subject statistical analysis on the muscle response onset times (in milliseconds) obtained by running the *detrended-integrated* signal analysis on 1000 bootstrapped data sets for each of the four cue conditions of the two experiments (see materials and methods). Note that the cue conditions outlined below the cue conditions represent only left target conditions for clarity

| **Target** | Top target | | | | Bottom target | | | |
|---|---|---|---|---|---|---|---|---|
| | Valid Valid | Valid Invalid | Invalid Valid | Invalid Invalid | Valid Valid | Valid Invalid | Invalid Valid | Invalid Invalid |
| **Cue condition** | 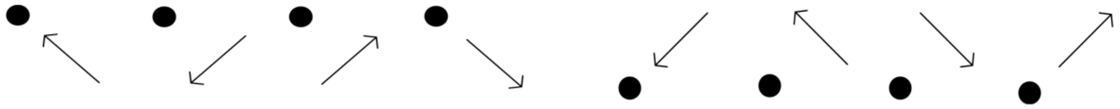 | | | | | | | |
| Experiment 1 Subjects | | | | | | | | |
| 1 | 81*,**,*** (80, 82) | 93+, ++ (90, 97) | 102 (98, 105) | 101 (98, 104) | 83**,*** (82, 85) | 86+, ++ (83, 88) | 106 (103, 110) | 100 (95, 105) |
| 2 | 82*,**,*** (81, 83) | 93+, ++ (90, 96) | 104 (100, 107) | 104 (100, 108) | 84**,*** (83, 85) | 86+, ++ (82, 90) | 107 (103, 111) | 100 (96, 104) |
| 3 | 90*,**,*** (88, 92) | 94+, ++ (90, 99) | 104 (99, 108) | 104 (100, 107) | 90*,**,*** (89, 92) | 94+, ++ (90, 97) | 100‡ (96, 104) | 105 (101, 109) |
| 4 | 88*,**,*** (86, 89) | 92+, ++ (88, 96) | 99 (96, 102) | 101 (96, 106) | 90*,**,*** (88, 91) | 95 ++ (90, 100) | 96‡ (92, 101) | 106 (101, 111) |
| 5 | 86**,***(84, 88) | 91+, ++ (87, 95) | 101‡ (97, 104) | 107 (103, 111) | 88**,*** (87, 89) | 85+, ++ (81, 89) | 106 (102, 110) | 102 (97, 106) |
| 6 | 82*,**,***(81, 83) | 89+, ++ (85, 93) | 102 (98, 106) | 106 (102, 110) | 84*,**,*** (82, 85) | 87+, ++ (84, 91) | 104 (99, 108) | 109 (104, 113) |
| 7 | 90*,**,*** (88, 91) | 95+, ++ (91, 98) | 102 (98, 105) | 103 (99, 108) | 88*,**,*** (86, 90) | 90+, ++ (87, 93) | 102 (99, 105) | 103 (99, 107) |
| 8 | 86*,**,***(84, 87) | 92+, ++ (88, 95) | 98 (95, 102) | 97 (92, 101) | 86*,**,*** (84, 87) | 94 (90, 97) | 97 (93, 102) | 97 (94, 101) |
| 9 | 86*,**,***(84, 87) | 91+, ++ (86, 96) | 110 (107, 113) | 101 (95, 108) | 90**,*** (88, 91) | 88+, ++ (84, 92) | 103 (99, 107) | 107 (103, 111) |
| 10 | 88*,**,***(87, 90) | 98++ (92, 103) | 101 (96, 106) | 105 (101, 109) | 90**,*** (89, 92) | 89+, ++ (85, 94) | 99‡ (96, 102) | 107 (104, 110) |
| 11 | 85*,**,***(84, 86) | 92+, ++ (89, 96) | 100 (96, 104) | 100 (95, 105) | 86**,*** (85, 88) | 90+, ++ (86, 94) | 98 (94, 102) | 101 (97, 105) |
| 12 | 85*,**,*** (83, 87) | 93+ (89, 97) | 100 (96, 104) | 99 (95, 103) | 85*,**,*** (84, 87) | 96 (90, 102) | 101 (97, 105) | 102 (97, 107) |
| Experiment 2 Subjects | | | | | | | | |
| 1 | 88**,*** (86, 90) | 89+, ++ (86, 92) | 105 (100, 110 ) | 103 (99, 107) | 90*,**,*** (88, 92) | 87+,++ (84, 89) | 104 (99, 114) | 107 (100, 114) |
| 2 | 94**,*** (92, 96) | 94+, ++ (91, 97) | 106 (100, 112) | 105 (99, 111) | 94**,*** (92, 96) | 93+,++ (90, 96) | 107 (104, 111) | 112 (106, 118) |
| 3 | 90**,*** (88, 92) | 91+, ++ (87, 94) | 100 (97, 103) | 103 (97, 109) | 90*,**,***(88, 92) | 86+,++ (83, 89) | 104 (99, 109) | 106 (100, 111) |
| 4 | 90**,*** (87, 93) | 91+, ++ (88, 93) | 105 (101, 109) | 106 (102, 110) | 91**,*** (90, 93) | 90+,++ (88, 93) | 107 (102, 112) | 103 (96, 110) |
| 5 | 89**,*** (87, 91) | 89+, ++ (86, 93) | 108 (102, 113) | 109 (103, 114) | 89**,*** (87, 91) | 91+,++ (88, 94) | 105 (100, 110) | 101 (96, 105) |
| 6 | 87**,*** (85, 89) | 88+, ++ (86, 91) | 102 (97, 107) | 103 (98, 108) | 87**,*** (85, 89) | 88+,++ (86, 91) | 104 (100, 108) | 105 (100, 110) |
| 7 | 91**,*** (89, 94) | 90+, ++ (87, 93) | 101 (95, 107) | 102 (96, 108) | 91**,*** (89, 93) | 93+,++ (90, 96) | 10 (97, 108) | 108 (103, 113) |
| 8 | 90**,*** (88, 92) | 91+, ++ (88, 94) | 102 (99, 106) | 103 (100, 107) | 89**,*** (86, 91) | 89+,++ (86, 92) | 103 (98, 107) | 104 (99, 108) |
| 9 | 88**,*** (86, 90) | 89+, ++ (86, 92) | 103 (97, 110) | 104 (98, 111) | 88**,*** (83, 87) | 87+,++ (84, 90) | 101 (95, 107) | 106 (99, 112) |
| 10 | 83*,**,*** (81, 85) | 86+, ++ (83, 89) | 104 (98, 109) | 105 (99, 110) | 85**,*** (83, 87) | 83+,++ (80, 87) | 102 (96, 108) | 97 (91, 102) |
| 11 | 88**,***(86, 91) | 89+, ++ (86, 91) | 102 (97, 106) | 103 (98, 107) | 87**,*** (84, 89) | 87+,++ (84, 90) | 107 (103, 112) | 110 (106, 114) |

First experiment's subjects 1, 4, 2, 8, 5, 7, 12 and 13 correspond to second experiment's subjects 1-5, 7, 10 and 11. The data are reported as mean (95% confidence interval). *, statistically significant difference between the Valid-Valid cue condition and the Valid-Invalid cue condition; **, statistically significant difference between the Valid-Valid cue condition and the Invalid-Valid cue condition; ***, statistically significant difference between the Valid-Valid cue condition and the Invalid-Invalid cue condition; +, statistically significant difference between the Valid-Invalid cue condition and the Invalid-Valid cue condition; ++, statistically significant difference between the Valid-Invalid cue condition and the Invalid-Invalid cue condition; ‡, statistically significant difference between the Invalid-Valid cue condition and the Invalid-Invalid cue condition.

### Supplementary Results: Experiment 2

*Between-methods contrasts*

For the second experiment, we compared the two methods of analysis on each of the eleven positive express responses producers (see materials and methods of the main manuscript for details), thus including the exemplar subject's data shown in figure 10 of the main manuscript. For this subject, the detrended-integrated signal analysis on the original data set showed that the express muscle response onset time in the Valid-Valid cue condition was 92ms for the top target and 93ms for the bottom target. These results were consistent with those obtained by running the detrended-integrated signal and ROC analyses on 1000 bootstrapped data sets (Supplementary Figure 1C and D; Supplementary Table 1: experiment 2, subject 4). Further, the single-subject analysis showed consistent (i.e. non-statistically significant, $p>0.05$) express response initiation times between the detrended-integrated signal and ROC analysis methods among the eleven express response producers of the second experiment (Supplementary Table 1), except for three subjects (Supplementary Table 1: experiment 2, subjects 5, 7 and 9).

*Between-cue conditions contrasts*

For the second experiment, we compared the express muscle response initiation between the four different cue conditions for each of the eleven positive express responses producers (see materials and methods of the main manuscript for details), thus including the exemplar subject's data shown in figure 10 of the main manuscript. For this subject, the detrended-integrated signal analysis on the original data set showed that the express muscle response onset time for both the top and bottom targets was ~93ms when the reaching direction was validly cued (i.e. Valid-Valid and Valid-Invalid cue conditions), and ~106ms when it was invalidly cued (i.e. Invalid-Valid and Invalid-Invalid cue conditions; see materials and methods of the main manuscript for details). These results were consistent with those observed by running the detrended-integrated signal analysis on 1000 bootstrapped data sets (Supplementary Figure 2C and D; Supplementary Table 2: experiment 2, subject 4). Consistently, the single-subject analysis showed that the express muscle response onset time of this subject was significantly earlier ($p<0.05$) when the reaching direction was validly (i.e. Valid-Valid and Valid-Invalid cue conditions) than invalidly cued (Supplementary Figure 2C and D; Supplementary Table 2: experiment 2, subject 4). The results of the exemplar subject were consistent among the eleven positive express response producers of the first experiment (Table 2: experiment 2).
